# Zebrahub-Multiome: Uncovering Gene Regulatory Network Dynamics During Zebrafish Embryogenesis

**DOI:** 10.1101/2024.10.18.618987

**Authors:** Yang Joon Kim, Shruthi Vijay Kumar, Benjamin Iovino, Alejandro A. Granados, Sarah Ancheta, Max Frank, Xiang Zhao, Kyle Awayan, Amanda Seng, Michael Borja, Sheryl Paul, Honey Mekonen, Ritwicq Arjyal, Angela Detweiler, Yasin Şenbabaoğlu, Rafael Gómez-Sjöberg, Norma Neff, Merlin Lange, Löıc A. Royer

## Abstract

During embryonic development, gene regulatory networks (GRNs) drive molecular differentiation of cell types. However, the temporal dynamics of these networks remain poorly understood. Here, we present Zebrahub-Multiome, a single-cell multiomic atlas that captures chromatin accessibility and gene expression from 94,562 cells across six stages of zebrafish embryogenesis (10-24 hours post-fertilization), capturing key developmental stages from the end of gastrulation to the onset of organogenesis. By measuring regulatory element activity alongside transcriptional output from the same cells, we identify 640,000 cis-regulatory elements organized into 402 hierarchically structured modules corresponding to specific developmental pathways. Early embryonic stages employ broadly shared regulatory programs that progressively fragment into lineage-specific modules. Time-resolved gene regulatory network inference reveals that transcription factors undergo functional transitions – from multilineage regulators to specialized, lineage-committed factors. These quantitative measurements reveal the regulatory network rewiring that drives cell fate specification. Our interactive web portal (zebrahub.org/epigenomics) enables exploration of gene dynamics, regulatory networks, and perturbation predictions, providing a quantitative framework for understanding vertebrate developmental regulation.

## Introduction

Vertebrate embryonic development transforms a single fertilized cell into a complex organism through precisely orchestrated waves of cell division, differentiation, and morphogenesis. This remarkable process is controlled by gene regulatory networks (GRNs) – interconnected systems where transcription factors (TFs) bind to *cis*-regulatory elements to control target gene expression in space and time^1,2^. Understanding how these regulatory networks dynamically reconfigure across developmental stages and emerging cell types remains a fundamental challenge in developmental biology^3^, with implications for regenerative medicine^4,5^, evolutionary biology^6,7^, and our understanding of developmental disorders^8,9^.

The past decade has witnessed a revolution in our ability to decode developmental programs at unprecedented resolution. Single-cell RNA sequencing (scRNA-seq) technologies have enabled comprehensive mapping of cell type emergence across entire developing organisms^10^. These advances have produced developmental cell atlases for major model organisms including *C. elegans*^11^, *Drosophila*^12^, mouse^13,14^, and zebrafish^15–20^ However, while these transcriptomic atlases reveal *what, when*, and *where* genes are expressed, they provide limited insight into *how* gene expression is regulated.

This regulatory layer has become accessible through singlecell multi-omic technologies that simultaneously profile chromatin accessibility and gene expression. By measuring open chromatin regions (via scATAC-seq^21^) alongside transcriptomes, these approaches directly identify active regulatory elements and their transcriptional effects during cell fate specification^22–25^. This dual measurement enables more accurate GRN inference by linking TF binding sites to target gene expression^26–29^.

Despite these advances, a critical dimension remains underexplored: the temporal dynamics of gene regulatory networks during development, i.e., how transcription factors and their target genes change over time or developmental lineages. Most single-cell studies capture a single static snapshot or focus on specific lineages^27,30^, missing how regulatory programs evolve continuously across developmental time. While some studies have attempted temporal analysis^31,32^, they typically rely on computational inference rather than direct temporal sampling. Furthermore, studies combining temporal resolution with multi-omic measurements have been limited to specific tissues or processes^33,34^, leaving whole-organism regulatory dynamics largely uncharted.

The zebrafish embryo provides an ideal system to address these challenges. Its external development, optical transparency, and rapid embryogenesis allow precise temporal sampling of all cell types simultaneously^35^. The period from 10 to 24 hours post-fertilization (hpf) encompasses critical developmental transitions: the completion of gastrulation, somitogenesis, and the onset of organogenesis, during which major cell lineages diverge and acquire specialized functions^35,36^.

Here, we present Zebrahub-Multiome, a comprehensive time-resolved single-cell multiomic atlas that captures the regulatory dynamics of zebrafish embryogenesis. By profiling chromatin accessibility and gene expression from the same cells at six developmental stages (10 to 24 hpf), we assembled data from over 94,000 cells to create an integrated view of how gene regulation unfolds across developmental time and cell types. Our analysis shows that chromatin accessibility is organized into a continuous hierarchy of regulatory modules corresponding to specific developmental processes and pathways. Through systematic annotation of roughly 640,000 regulatory elements clustered into 402 peak modules, we identified pathway-specific regulatory programs and extracted modulespecific regulatory sub-networks. This approach revealed three key principles: (i) early embryonic stages share broad regulatory programs that progressively commit into lineage-specific modules, (ii) individual transcription factors transition from multi-lineage to specialized roles as development advances, and (iii) chromatin accessibility and gene expression exhibit complex temporal relationships that vary by genomic context, with promoters providing stable foundations while distal elements confer cell-type specificity.

To enable the functional interpretation of these regulatory dynamics, we integrated all datasets and interactive analyses into a web portal, Zebrahub-Multiome, which includes our peak-derived modules with GRN inference and *in silico* genetic perturbation analysis, predicting how transcription factor knockout affects development.

## Results

### A time-resolved single-cell multi-omic atlas of zebrafish development

To investigate the epigenomic and transcriptomic profiles during cell fate specification in zebrafish embryogenesis, we conducted a single-cell Multiome assay (see Methods), which simultaneously captures gene expression (RNA) and chromatin accessibility (ATAC) from the same nuclei. We sampled six time points between 10 and 24 hours post-fertilization (hpf), a critical developmental window spanning from the end of gastrulation to the onset of organogenesis (Fig.1a; see Methods and Supp. Fig. S1 for the experimental protocol and data processing workflow) ^35^. After quality control, the dataset comprised 94,562 cells, with a median of 1,400 detected genes, 3,700 UMI counts, and 15,000 peaks per cell (Supp. Fig. S2).

To visualize the epigenomic and transcriptomic profiles of these cells, we first integrated data across timepoints and replicates using reciprocal PCA and LSI (see Methods; Fig. S1), then computed uniform manifold approximation and (UMAP) (Fig. 1a). We generated three embeddings: (i) RNA, (ii) ATAC, and (iii) Joint (a weighted nearest neighbor from RNA and ATAC modalities) ^37^, as shown in Fig. 1b, c, and d, Supp. Video 1, *this* online visualization, and available for exploration online in Zebrahub-Multiome.

**Figure 1:**
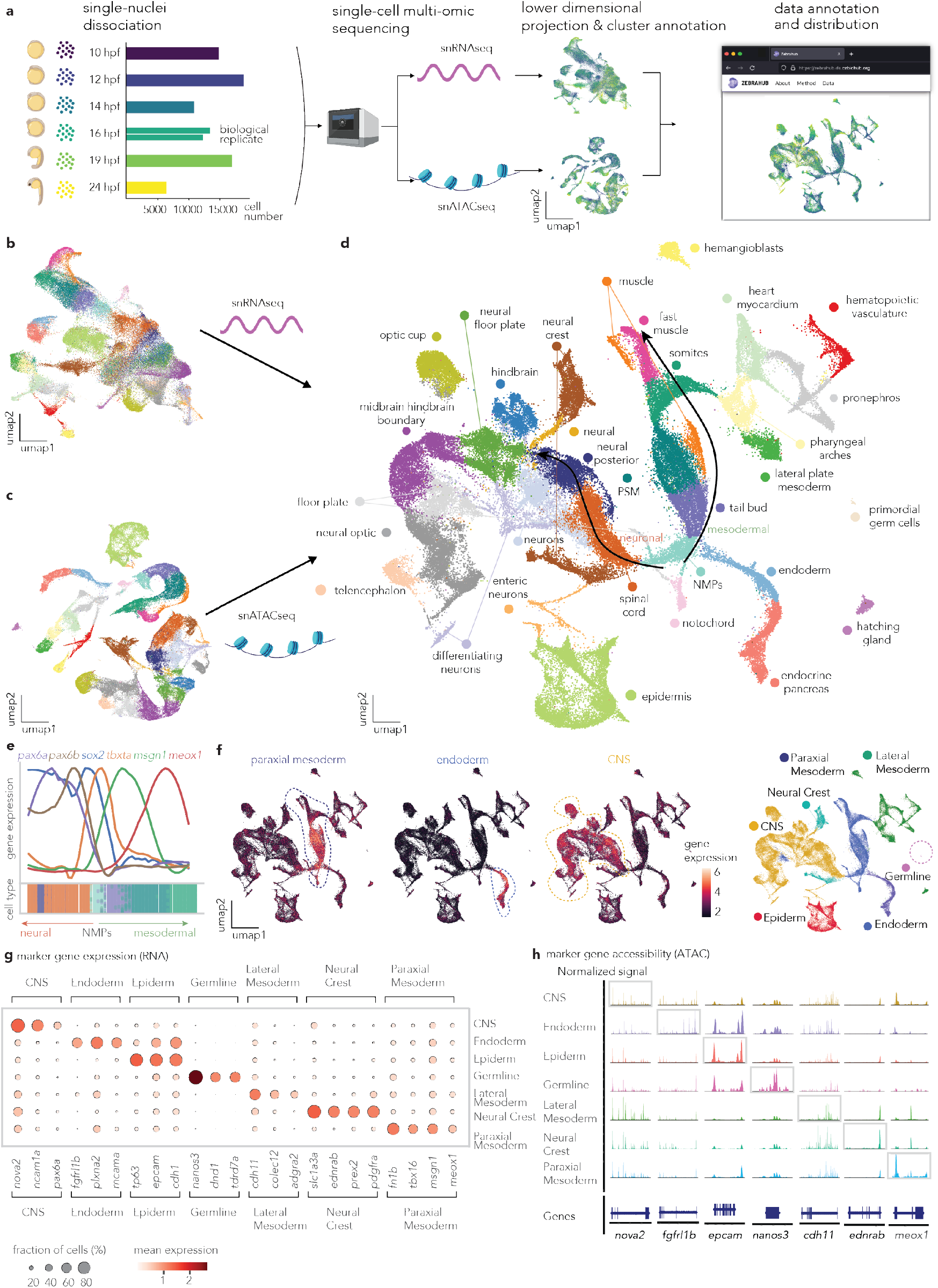
Building a time-resolved single-cell multi-omic atlas of zebrafish embryonic development. **a**, A schematic of the experimental and computational analysis workflow. From left to right, we show: First, the number of cells per time point and replicates. Second, single-nuclei dissociation from pooled zebrafish embryos at 10, 12, 14, 16, 19, and 24 hours post fertilization (hpf), followed by single-cell Multiome assay for both snRNA-seq and snATAC-seq. Third, computational analysis of single-cell multiome datasets, including dimensionality reduction, clustering, and cell type annotation. Lastly, a web portal allows the exploration of single-cell Multiome datasets online. **b-d**, UMAP representation of **b**, RNA modality, **c**, ATAC modality, and **d**, joint (RNA+ATAC) colored by annotated cell types, integrated over all timepoints. **e**, Gene expression (RNA) dynamics along the pseudotemporal axis for neural (left) and mesodermal (right) trajectories from the neuro-mesodermal progenitors (NMPs). Gene expression is normalized to emphasize their dynamics. **f**, (Left) A joint UMAP representation of gene expression patterns for major tissues: mesoderm, endoderm, and neuro-ectoderm using marker genes (See Methods for the list of marker genes for each tissue or lineage), and (Right) colored by tissue annotation. (See Table 1 for the dictionary mapping of cell types to tissues.) **g**, A dot plot showing marker gene expression (RNA) across major tissue categories shown in **f**. Circle size represents the fraction of cells expressing each gene, and color intensity indicates mean expression level within each tissue. **h**, Marker gene accessibility profiles (ATAC) across tissue categories shown in **f**. Genome tracks show normalized chromatin accessibility signals for representative marker genes.

Our first question was whether multimodal integration improves cell type annotation beyond what is possible by individual modalities. We performed cross-modality validation using neighborhood purity and trajectory conservation analyses (see Supp. Fig. S3 and Methods). Neighborhood purity quantifies whether cells sharing similar molecular profiles also share cell type identity – high purity indicates that a modality effectively groups cells of the same type together while separating different types (Supp. Fig. S3b). Consistent with this metric, ATAC-based neighborhoods showed higher purity than RNA-based neighborhoods for unsupervised leiden clusters from each modality (Supp. Fig. S3d), confirming ATAC’s superior ability to delineate discrete cellular identities.

**Table 1:** Cell type to lineage mapping for coarse-grained analysis. Individual cell types from the original Zebrahub annotation were grouped into seven major developmental lineages based on their embryological origins and shared developmental programs.

| Lineage | Cell Types |
| --- | --- |
| CNS | Neural, neural optic, neural posterior, neural telencephalon, neurons, hind-brain, midbrain-hindbrain boundary, optic cup, spinal cord, differentiating neurons, floor plate, neural floor plate, enteric neurons |
| Neural Crest | Neural crest |
| Paraxial Mesoderm | Somites, fast muscle, muscle, PSM (presomitic mesoderm), NMPs (neuromesodermal progenitors), tail bud, notochord |
| Lateral Mesoderm | Lateral plate mesoderm, heart myocardium, hematopoietic vasculature, pharyngeal arches, pronephros, hemangioblasts, hatching gland |
| Endoderm | Endoderm, endocrine pancreas |
| Epiderm | Epidermis |
| Germline | Primordial germ cells |

However, high neighborhood purity alone does not guarantee accurate representation of continuous developmental relationships, where cell types diverge from shared ancestors along progressive trajectories. We assessed whether each modality preserves these differentiation progressions by assessing ranked correlations between unsupervised pseudotime estimates and expected cell type progressions (Supp. Fig. S3e). RNA-based neighborhoods maintained developmental trajectories for both mesodermal and neuro-ectodermal lineages well (conservation scores = 1.0 and 0.7, respectively), while ATAC-based neighborhoods showed substantially reduced conservation (scores = 0.4 and −0.5, respectively; Supp. Fig. S3e). This striking difference demonstrates that RNA expression more faithfully captures the continuous nature of cellular differentiation during embryogenesis. The joint embedding successfully integrated these complementary strengths – achieving both high neighborhood purity for discrete cell type identification and accurate trajectory conservation for developmental continuity. Thus, multimodal integration of both ATAC and RNA data captures the dual nature of development, i.e., discrete cell type identities existing within continuous differentiation trajectories, providing a more complete representation than either modality alone.

Given these complementary strengths, we performed cell type annotation using the joint UMAP embedding, which optimally balances cell type separation and developmental continuity. Cell type identities were assigned through RNA-based label transfer from a reference atlas^20^ followed by expert curation using marker gene expression and zebrafish ontology terms (see Methods)

Next, we examined whether our integrated atlas correctly captures well-known developmental gene expression patterns and molecular signatures. Gene expression along pseudotemporal trajectories from neuro-mesodermal progenitors (NMPs) through their neural and mesodermal derivatives revealed sequential gene activation matching the developmental sequence shown by Lange et al.^20^ (Fig. 1e). Furthermore, marker gene expression patterns strongly correlated with their corresponding chromatin accessibility profiles across all major lineages (Fig. 1f-h; see Methods and Table 1 for complete marker gene lists and tissue annotation details), confirming that our cell type annotations reflect genuine molecular states^38^.

Having generated a time-resolved, multi-omic developmental cell atlas during zebrafish embryogenesis, we recognized that integrating chromatin accessibility and gene expression data provides a unique opportunity to capture the temporal sequence of gene regulation, where chromatin opening events precede and enable transcriptional activation. To systematically exploit this multi-omic resolution and understand how regulatory programs unfold across the developmental landscape, we next investigated how individual genes exhibit coordinated dynamics across both cell types and developmental stages.

### Gene regulation dynamics reveal temporal coupling of chromatin accessibility and transcription across cell types and developmental stages

For genes to be transcribed, the relevant genomic regions, promoters, and enhancers must first become accessible to transcription factors^39^. While chromatin accessibility changes typically precede gene expression changes^24,40^, the temporal dynamics of this relationship during development remain poorly understood. To address this gap, we developed a genelevel dynamics analysis that constructs a regulatory activity profile for each gene across developmental stages and cell types. This approach enables the identification of genes with similar dynamics to test whether such similarity reflects shared biological functions and developmental roles.

To implement this approach, we first profiled the chromatin accessibility for each gene using Signac^41^, resulting cells-bygene activity matrix, where gene activity represents the sum of chromatin accessibility over the gene body for each gene. To mitigate the sparsity inherent in single-cell ATAC-seq and RNA-seq datasets, we employed a metacell aggregation strategy^42^ (Fig. 2a), then computed mean gene activity (ATAC) and expression (RNA) values across cell types at each developmental stage to create the gene dynamics vectors (Fig. 2b; Fig. S5; see Methods). For downstream analyses, we selected 5,068 highly variable genes (a union of 4000 highly variable genes from either RNA or ATAC modality, respectively).

**Figure 2:**
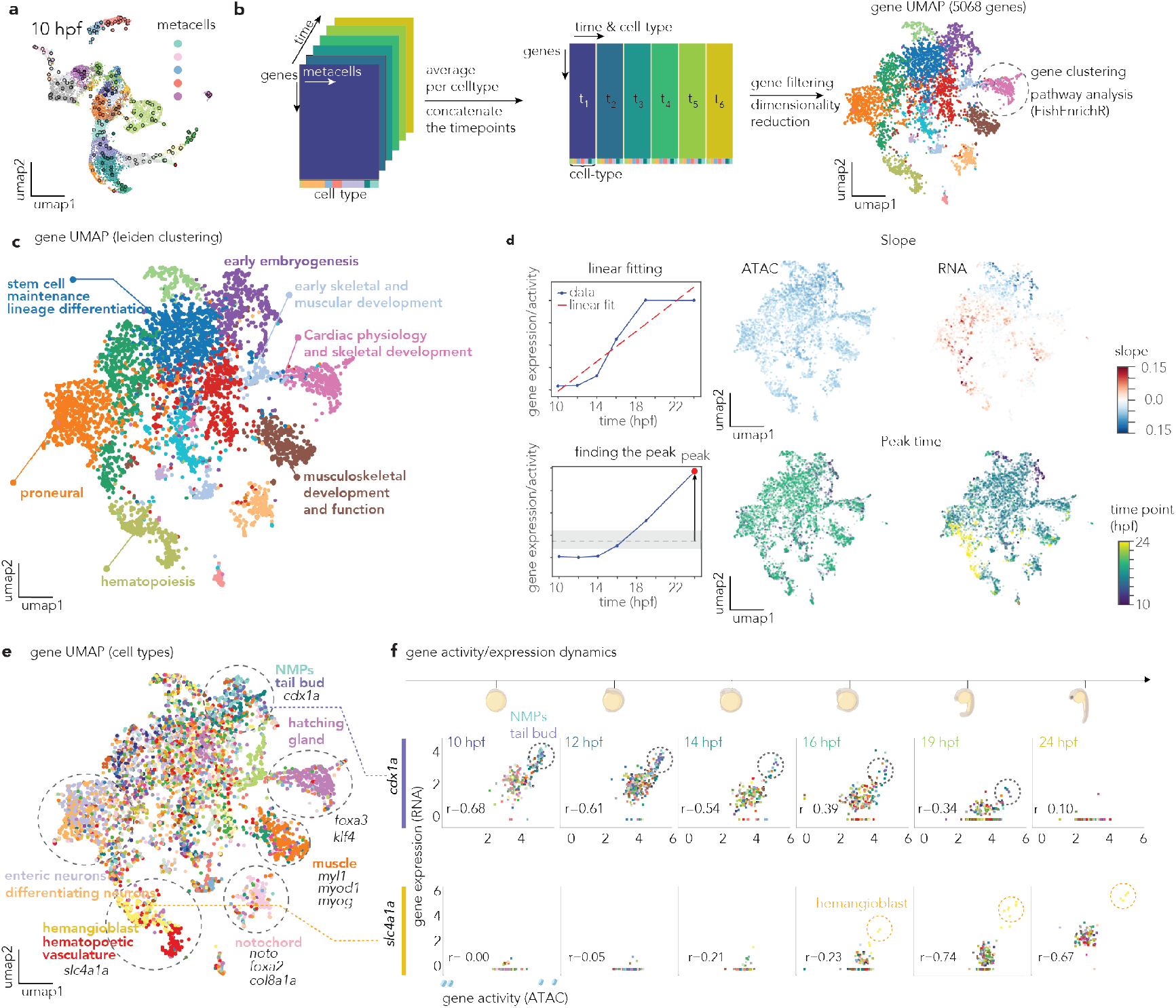
Gene-centric analysis reveals distinct temporal dynamics of chromatin accessibility and gene expression during zebrafish embryogenesis. **a**, A UMAP representation of joint modalities (RNA+ATAC) at 10 hpf showing single-cells with metacells colored by the cell type annotation in Fig. 1d. **b**, Computational workflow for gene dynamics analysis. Metacells are computed from single-cell ATAC-seq data, then averaged by cell type and time point to create gene expression and accessibility vectors. Each gene is represented as a vector capturing its dynamics across all cell types and developmental stages. After gene filtering, clustering, and pathway analysis, genes are dimensionally reduced using UMAP to visualize functionally related gene modules. **c**, Gene UMAP (5,068 genes), where each point represents a gene, colored by Leiden clustering. Genes cluster based on their shared dynamics across cell types and developmental stages. Major developmental processes are annotated, including early embryogenesis, stem cell maintenance, cardiac physiology, musculoskeletal development, and hematopoiesis genes. **d**, Temporal pattern analysis of gene dynamics. Left: Linear fitting approach to quantify gene expression/activity slopes over time. Right: Gene UMAP colored by slope values for ATAC (left) and RNA (right), and peak expression timing (bottom), revealing genes with increasing, decreasing, or peaked temporal patterns. **e**, Gene UMAP colored by the cell type where each gene shows the highest expression, demonstrating cell-type-specific gene modules within the dynamics-based clustering. **f**, Examples of gene activity/expression dynamics over developmental time. Top: *cdx1a* shows decreasing correlation between ATAC and RNA over time in NMP/tail bud populations. Bottom: *slc4a1a* shows increasing correlation over time in hemangioblast populations. Each time point shows scatter plots with Pearson correlation coefficients, colored by the predominant cell type.

Using this matrix of gene dynamics, we analyzed how genes cluster based on their shared temporal and cell-type-specific regulatory patterns, and visualized the data using UMAP, followed by leiden clustering (resolution=0.7) (Fig. 2b, See Methods). These unsupervised clusters corresponded to functionally coherent developmental processes. Gene set enrichment analysis using FishEnrichr^43,44^ identified distinct modules including “early embryogenesis,” “stem cell maintenance and lineage differentiation,” “cardiac physiology and skeletal development,” “musculoskeletal development and function,” “hematopoiesis,” and “proneural genes,” validating that genes with coordinated regulatory dynamics reflect shared biological functions (Fig. 2c; Supp. Fig. S6).

Next, we quantified the overall temporal behavior of individual genes. We first averaged gene activity and expression values, respectively, across all cell types for each developmental timepoint, then performed linear regression to calculate the slope of these averaged values over developmental time (Fig. 2d). This approach captured the overall temporal trajectory of each gene’s regulatory activity from 10 hpf to 24 hpf. The analysis revealed distinct temporal patterns between chromatin accessibility and gene expression. Chromatin accessibility (ATAC) showed predominantly mild but consistent decreasing trends across most genes, with peak accessibility occurring around 14-16 hpf (Fig. 2d, top left). This suggests that chromatin is generally more open during early-to-mid developmental stages and gradually becomes more restricted toward 24 hpf, reflecting the progressive commitment of cell fates during development. In contrast, gene expression (RNA) exhibited a broader dynamic range with both increasing and decreasing temporal patterns, indicating more diverse dynamics for transcriptional activation across developmental processes (Fig. 2d, top right). When we visualized these slope values on the gene UMAP, functionally related genes often exhibited similar temporal dynamics (Fig. 2d). Early developmental genes predominantly showed decreasing trends, while genes involved in later differentiation processes showed increasing patterns, demonstrating that temporal regulatory behavior reflects functional roles in development (Fig. 2d). To examine cell-type specificity of gene expression, we visualized each gene colored by the cell type where it exhibits maximum expression, revealing distinct cell-type-specific gene modules organized by their dynamic regulatory patterns (Fig. 2e).

These temporal dynamics are exemplified by two genes with contrasting regulatory patterns (Fig. 2f). *cdx1a*, a homeobox gene involved in anterior-posterior axis formation^45^, shows decreasing levels for both chromatin accessibility and gene expression over time, particularly in NMP and tail bud populations. This pattern reflects the early activation of this developmental regulator, where chromatin remains accessible even as expression declines with developmental progression^46^. In contrast, *slc4a1a*, involved in hematopoiesis and erythrocyte differentiation^47^, exhibits increasing patterns over time in hemangioblast populations. Here, chromatin accessibility precedes transcriptional activation, illustrating the classic model where regulatory element opening enables subsequent gene expression during blood cell differentiation^48^.

These gene-level dynamics analyses reveal that regulatory coupling between chromatin accessibility and gene expression follows distinct temporal patterns, reflecting the functional roles of genes in developmental lineage specification rather than uniform relationships across development. The complete analysis is available online as an interactive resource for all genes used for Figure. 2.

### Chromatin accessibility profiles reveal shared and unique epigenetic regulatory programs

While our analysis of gene dynamics revealed coordinated transcriptional regulation throughout development, this approach analyzed chromatin accessibility only within regions of the gene body and their immediate proximal promoters. To capture the full regulatory landscape, including both proximal and distal regulatory elements such as enhancers and silencers that operate over long genomic distances, we expanded our analysis to examine discrete regulatory regions throughout the genome. We developed a peak-centric approach that analyzes individual ATAC-seq peaks – discrete regions of open chromatin representing potentially active regulatory elements – across cell types and developmental stages (Fig. 3a). We analyzed all 640,830 ATAC-seq peaks using pseudobulk profiles aggregated across 32 cell type clusters and 6 developmental time points, creating a 190-dimensional accessibility profile for each regulatory element (Fig. 3b; Supp. Fig. S8).

**Figure 3:**
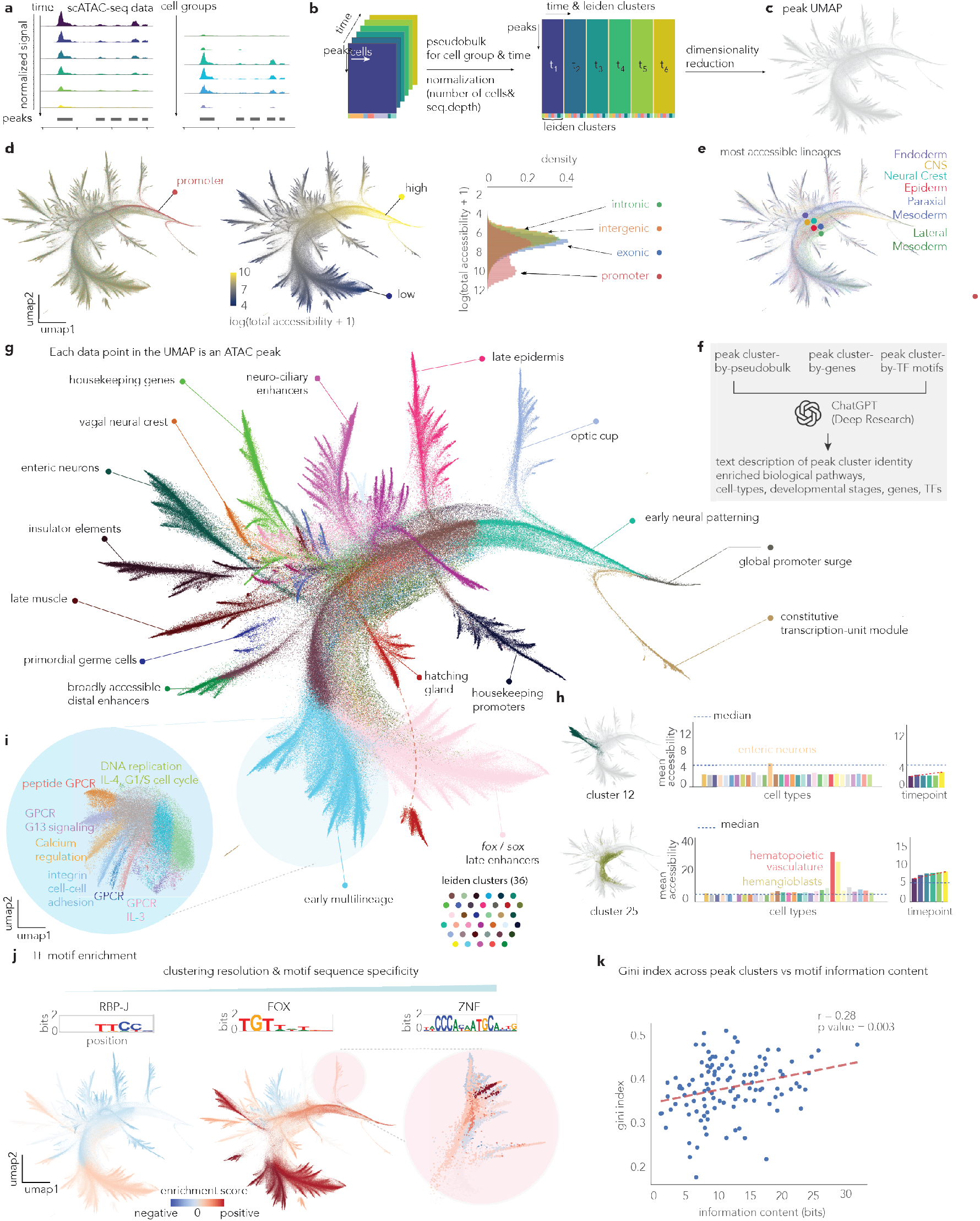
Peak-centric analysis reveals regulatory programs organized by developmental pathways and upstream transcription factors. **a**, Schematic of scATAC-seq data analysis workflow capturing variability of chromatin accessibility across developmental stages and cell types. **b**, Computational pipeline for peak dynamics analysis. Peaks are grouped by cell types and time points, converted to pseudobulk profiles, normalized, and subjected to dimensionality reduction to generate a peak UMAP for visualization. **c**, UMAP visualization of peak accessibility patterns showing the overall structure of regulatory element clustering. **d**, Peak characteristics analysis showing distribution by (left) genomic annotation (exonic, intergenic, intronic, promoter) and (center) chromatin accessibility levels, and (right) the accessibility levels for each peak type. **e**, UMAP highlighting the most accessible lineages as defined in Table.1. **f**, Large Language Model (LLM) based annotation of the peak clusters. The coarse (36 clusters) and fine (402 clusters) Leiden clusters were systematically annotated using ChatGPT, based on their accessibility across cell types and time points, the list of associated genes, and enriched transcription factors, to identify enriched biological pathways and processes. **g**, Peak UMAP colored by coarse Leiden clusters (36 clusters), highlighting several clusters with annotations from **f. h**, Averaged chromatin accessibility of two coarse Leiden clusters across cell types or time points, representing their cell-type and time-point specificity. **i**, Sub-clustering of a coarse cluster colored by the fine Leiden clusters and annotation from **f. j**, Transcription factor motif enrichment analysis across different clustering resolutions, showing representative motifs with varying sequence specificity (RBP-J, FOX, and ZNF transcription factor motifs). **k**, A scatter plot showing the gini index of transcription factor motif enrichment computed across the peak clusters and the information content for sequence motif. Pearson correlation coefficient and linear fit are shown.

Dimensionality reduction of this complex dataset revealed a hierarchical tree-like structure (Fig. 3c; see Methods), particularly striking when visualized in three dimensions (see Supp. Video 2 and *this* online visualization). This hierarchical organization persists when using different hyperparameters (Supp. Fig.S9), and different dimensionality reduction approaches (Supp. Fig.S10), yet largely disappears when analyzing only the subset of highly variable peaks (Supp. Fig. S11b).

The robustness of this structure across various methods indicates that the hierarchy reflects a fundamental organization of the regulatory genome. The tree’s root consists of promoter-proximal peaks exhibiting consistently high accessibility across all cell types and developmental stages (Fig. 3d), consistent with the well-established principle that promoters maintain constitutive accessibility while regulatory specificity is conferred by distal enhancers^49–53^. In contrast, the remaining regulatory elements – comprising the vast majority of peaks (intergenic 36.4%, intronic 40.6%, exonic 18.9% vs. promoter 4.1%) – are scattered without type-specific clustering (Fig. 3d; Supp. Fig. S11c). The tree’s trunk reveals parallel layers (Fig. 3e) corresponding to major developmental lineages: paraxial and lateral mesoderm, epiderm, endoderm, CNS, and neural crest (Table 1). These primary branches progressively subdivide into tens of smaller branchlets (Fig. 3g,Supp. Fig. S14; Supp. Video 2; interactive visualization available online).

To dissect this hierarchical organization, we applied twolevel Leiden clustering, identifying 36 major clusters and 402 fine-grained sub-clusters (Supp. Fig. **??**a). Because 77% of peaks reside in functionally uncharacterized intergenic and intronic regions (Fig. 3d), we developed a systematic annotation approach using Large Language Models^**?**^. Each cluster was characterized based on three comprehensive data matrices as shown in Figure. 3f: (i) accessibility profiles across cell types and time points (Fig. 3h), (ii) associated target genes (Supp. Fig. S12), and (iii) enriched transcription factor motifs (Supp. Fig. S13). This multi-dimensional characterization enabled functional annotation of each regulatory module – 36 coarse clusters and 402 fine clusters in total (see Methods).

The annotations revealed that coarse clusters correspond to major developmental programs (late muscle development, enteric neuron formation, epidermis specification; Fig. 3g,h), while fine-grained sub-clusters capture more specific programs driven by specific signaling pathways and regulatory circuits (GPCR signaling, BMP/Wnt pathways, neural crest migration, cell cycle regulation; Fig. 3i; Supp. Fig. S14). Supporting this functional organization, each cluster displayed distinctive combinations of transcription factor motifs (3-15 motifs per cluster; Fig. 3j; Supp. Fig. S13), with a mild, but significant correlation between clustering resolution and motif specificity (r = 0.28, p = 0.003; Fig. 3k).

This hierarchical organization of regulatory elements, ranging from broadly accessible promoters to developmental pathway-specific enhancer modules, is potentially driven by the differential enrichment of transcription factor motifs, providing the genomic architecture for precise spatiotemporal control of gene expression during development. Having mapped this regulatory landscape, we next investigated how transcription factors coordinate these elements into functional gene regulatory networks.

### Cell-type- and stage-specific gene regulatory networks reveal conserved and dynamic gene regulatory modules

To understand how transcription factors coordinate the accessibility profiles of peak clusters, identified in our peak analysis, we constructed gene regulatory networks (GRNs) that map the relationships between TFs, *cis*-regulatory elements, and target genes, then examined how these networks are dynamically rewired across developmental stages and cell lineages (Fig. 4a).

**Figure 4:**
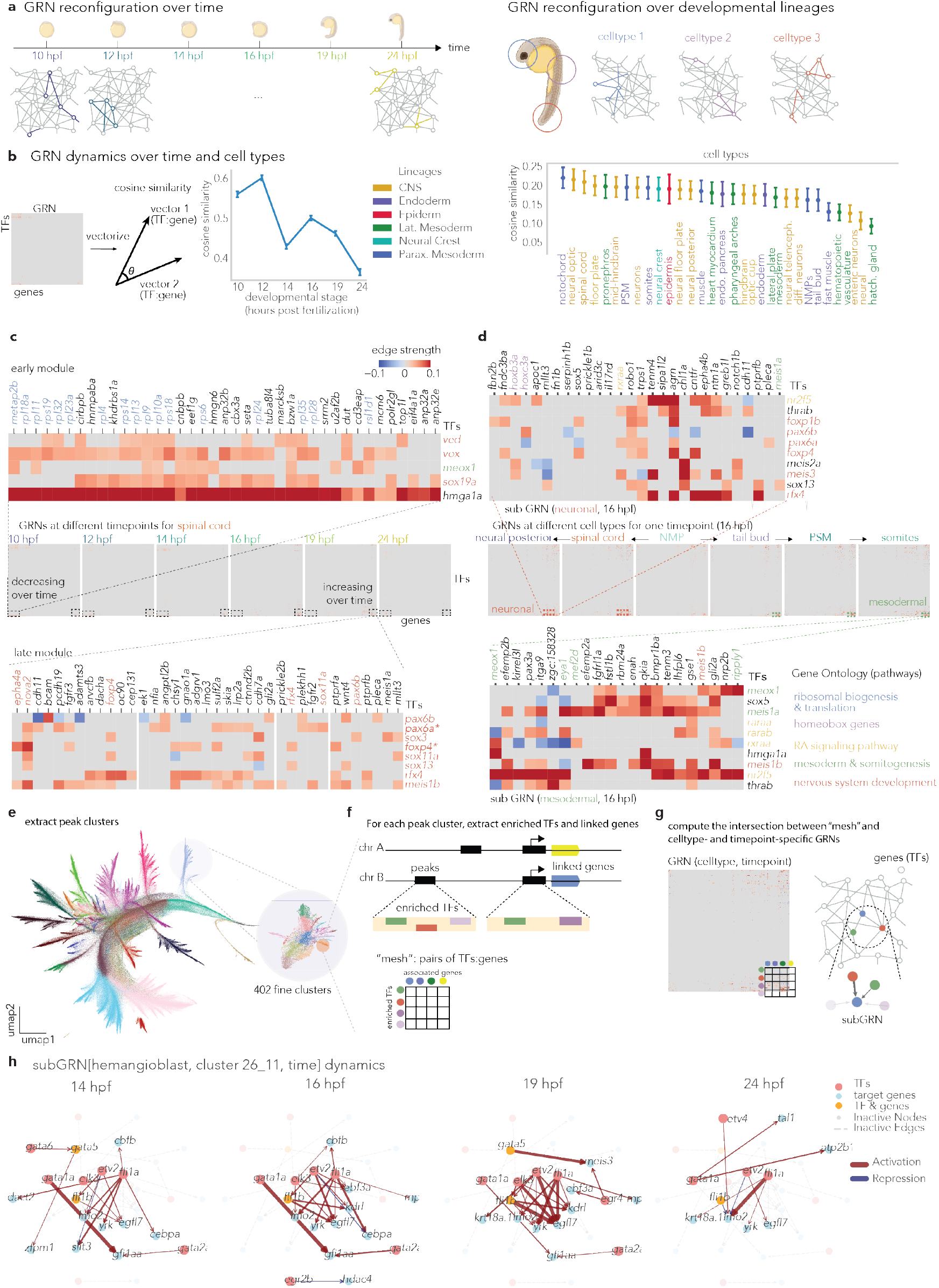
Dynamic Gene Regulatory Networks (GRNs) across developmental stages and cell types during zebrafish embryogenesis. **a**, A schematic representation of GRN reconfiguration (1) over developmental time and (2) across different cell types. **b**, GRN similarity quantification using cosine similarity. (Left) GRN similarity between cell types over developmental stages. (Right) GRN similarity between time points over cell types. The cell types were colored using “lineages” defined in Table 1. **c**, (Center) GRNs for spinal cord development across different developmental stages (10 to 24 hpf). Heatmaps display the edge strengths between transcription factors (TFs) and target genes, with both decreasing and increasing patterns highlighted. (Top, Bottom) Detailed view of the GRN heatmaps for spinal cord, showing specific TF-gene interactions for modules that are active in early (Top) and late (Bottom) time points. Edge strength is represented by color intensity, with red indicating positive and blue indicating negative regulation. **d**, GRNs for different cell types at the 16 hpf stage, including Neural Posterior, Spinal Cord, Neuromesodermal Progenitors (NMP), Tail Bud, Presomitic Mesoderm (PSM), and Somites.(Top, Bottom) Sub-GRNs of the 16 hpf stage GRN for neuronal (Top) and mesodermal (Bottom) lineages. Gene Ontology pathways associated with specific modules are annotated. Notable TFs and their target genes are highlighted, demonstrating lineage-specific regulatory patterns (also shown in c). **e**, Peak UMAP highlighted with coarse-grained Leiden clusters, and zoomed into a specific fine-grained Leiden cluster (optic cup). **f**, A scheme to create a mapping of transcription factors whose motifs are enriched in the peak cluster, and linked genes whose gene expression is highly correlated with peaks, resulting in a “mesh” of transcription factors-by-target genes. **g**, A scheme to extract the subGRN from a celltype-, and timepoint-specific GRNs using “mesh”. **h**, An example subGRN from neural floor plate over time, subsetted by regulatory program from the peak cluster “26 − 11”. Transcription factors are colored in red, target genes are colored in blue, and the transcription factors that are also target genes are colored in yellow. Inactive edges are colored grey to indicate the transient interactions that turn on or off between time points.

**Figure 5:**
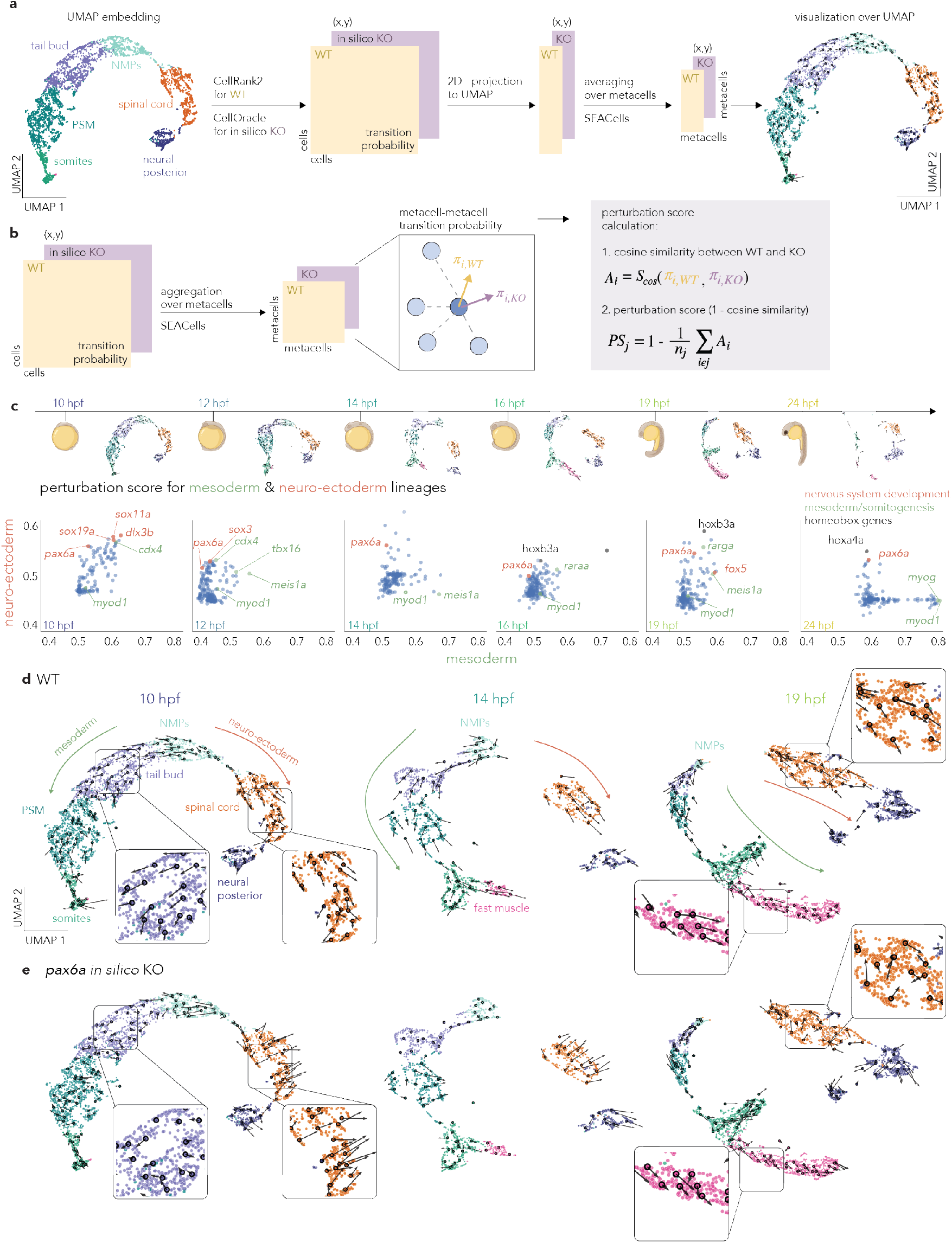
Cell-cell transition probability analysis and *in silico* perturbation using CellOracle. **a**, Workflow for computing and visualizing cell-cell transition probabilities. Starting from a 2D UMAP embedding, transition probabilities are calculated for wildtype (WT) and knock-out (KO) conditions, projected onto UMAP, and averaged over metacells for visualization. **b**, Schematic of metacell-metacell transition probability calculation and perturbation score computation using cosine similarity between WT and KO transition probabilities. **c**, Perturbation scores for mesoderm/neuro-ectoderm lineages across developmental time points (10 to 24 hpf). Each dot (blue) represents a transcription factor, with higher scores indicating stronger perturbation effects for either mesoderm (x-axis, green), or neuro-ectoderm (y-axis, red) lineages. **d**, UMAP visualizations of WT zebrafish embryo development from 10 to 24 hpf, with major cell types (dot color) and lineages (arrows with green for mesoderm, and red for neuro-ectoderm) annotated. Black arrows indicate predicted cell state transitions. **e**, UMAP visualizations of *in silico pax6a* knockout at 10, 14, and 19 hpf, showing altered cell state transitions compared to WT.

We used the CellOracle framework^27^ to compute time- and cell-type-specific GRNs by integrating scATAC-seq and scRNA-seq data (see Methods; Supp. Fig. S16). This approach models gene expression as a function of TF activity and regulatory element accessibility, capturing both developmental stage- and cell-type-specific regulatory programs (Fig. 4a).

To test whether regulatory networks reflect the progressive specialization of cell fates during development, we quantified GRN variability across cell types and time. First, we computed pairwise cosine similarities between all cell-type-specific GRNs at each developmental stage (Fig. 4b, left, See Methods for details). Consistent with the transition from pluripotent to specialized cellular states, mean GRN similarity decreased systematically from 0.56 at 10 hpf to 0.35 at 24 hpf (n = 903 cell type pairs per timepoint) – a 37% reduction in regulatory network similarity during somitogenesis. Second, we assessed temporal similarity by comparing GRNs from the same cell type across different timepoints (Fig. 4b, right). We found a low mean similarity of 0.20-0.25 within individual cell types, with comparable similarity across all developmental lineages (CNS, Endoderm, Epiderm, Germline, Lateral Mesoderm, Neural Crest, and Paraxial Mesoderm). Overall, these results suggest that temporal progression has a slightly stronger impact on regulatory network rewiring than cell-type differentiation during embryogenesis.

Next, to investigate cell-type-specific regulatory networks, we examined the spinal cord, a critical vertebrate structure^54^. At 24 hpf, spinal cord GRNs revealed substantially clearer regulatory relationships than whole-embryo networks (Fig. 4c; Fig. S17b and c), demonstrating that cell-type resolution is essential for capturing biologically meaningful regulatory interactions. Moreover, our analysis (Fig. 4c) shows that dynamic gene regulatory modules change over time, reflecting the shifting priorities of the developing embryo. Indeed, a module active in earlier stages (10 hpf), driven by the chromatin modifier *hmga1a*^55^, was responsible for activating genes encoding ribosomal subunits (Fig. 4c-top). This module was prominent in other cell-type-specific GRNs at early time points (Supp. Fig. S17e). In contrast, later-stage modules were enriched with transcription factors specific to neuronal development, as identified through gene ontology curation from ZFIN^54^ (Fig. 4c-bottom). The early activation of modules related to ribosomal biogenesis and translation suggests that the embryo prioritizes building cellular machinery to support cell cycle progression and protein production, crucial for cell fate specification. The involvement of *hmga1a* in regulating these early processes highlights the role of chromatin modifiers in orchestrating early developmental events^55^ (Fig. 4c-top).

To further explore cell-type-specific gene regulatory modules, we examined GRN dynamics across the differentiation of two major posterior lineages: neuro-ectoderm and paraxial mesoderm, both originating from the bipotent Neuro-Mesodermal Progenitors (NMPs^56^) (Fig. 4d). These progenitors can differentiate into either neuro-ectoderm (spinal cord and neural posterior) or paraxial mesoderm (presomitic mesoderm, somites, and fast muscle)^20^. At the early stage (10 hpf), the GRN architecture was highly similar across these developmental lineages, suggesting that the regulatory network structure is largely shared during the initial phase of development (Supp. Fig. S17e). By 16 hpf, we observed distinct GRN modules activated during the differentiation of each lineage, with neural-specific modules highlighted in red and mesodermal-specific modules in green (Fig. 4d). Further analysis of the genes within these modules (Fig. 4d-top and bottom) revealed that they are crucial for neural and mesodermal development, respectively^20,56^.

Overall, our results show that in early development, GRNs are largely conserved across cell types (Fig. 4b and Supp. Fig. S17e). However, as development progresses, these shared regulatory modules give way to lineage-specific gene regulatory programs (Fig. 4d), illustrating the shift from a common regulatory architecture to specialized modules over time.

### Dissection of gene regulatory networks using peak-derived regulatory programs

While cell-type and timepoint-specific GRNs provide detailed regulatory landscapes, their complexity (*>*2,000 TF-gene interactions per network) still obscures the identification of specific regulatory modules that drive distinct developmental programs within each cellular context. To identify functionally coherent subnetworks, we developed an approach that leverages the regulatory programs identified in our peak analysis to extract biologically disentangled GRN modules (Fig. 4e-g; Supp. Fig. S19).

Our rationale was that peak clusters represent co-accessible regulatory elements across cell types and developmental stages, indicating they operate under shared regulatory control (Fig. 4e). These co-accessible peak clusters should therefore be regulated by similar transcription factor combinations and control related target genes. We exploited this principle by creating “regulatory program meshes” – binary matrices that map transcription factors to target genes based on peak cluster organization (Fig. 4f). For each of the 402 finegrained peak clusters, we identified: (1) transcription factors whose binding motifs were significantly enriched within the cluster, and (2) genes whose expression was highly correlated with peak accessibility, resulting in binarized “meshes” for each peak cluster (Fig. 4f; See Methods for details). This process generated cluster-specific TF-by-gene matrices that serve as filters to extract relevant subGRNs from the cell-type and timepoint-specific networks generated by CellOracle (Fig. 4g).

This approach successfully identified modular regulatory programs with distinct temporal dynamics. For example, analysis of the regulatory program “26-11” in hemangioblasts revealed a 28-node subGRN comprising 11 transcription factors and 19 target genes (Fig. 4h). The network captured the well-characterized hemangioblast specification cascade, with the ETS family master regulatory Etv2^57,58^ and its downstream targets *fli1a, fli1b*, and *gata2a*^59,60^ *activating at 14 hpf, coinciding with hemangioblast emergence in posterior lateral plate mesoderm*^*61*^. *Network complexity peaked at 16-19 hpf as the GATA cascade expanded (gata1a,gata5,gata6*) and hematopoietic specification genes (*tal1, lmo2*)^62^ became active. By 24 hpf, the network consolidated to 9 edges as primitive hematopoiesis transitioned toward definitive programming^63,64^. Notably, several nodes functioned as both transcription factors and target genes (highlighted in yellow), indicating the presence of hierarchical regulatory cascades within the module. This regulatory program-based approach reduced network complexity by 99% while disentangling biologically meaningful modules, enabling detailed analysis of how specific regulatory circuits rewire during development.

Our comprehensive GRN analysis revealed two key principles governing the dynamics of regulatory networks during zebrafish development. First, regulatory networks progressively diversify from shared early programs to specialized cell-typespecific architectures (Fig. 4b-e; Fig. S17e,f). Second, by extracting regulatory program-based subGRNs from peak clusters, we identified discrete modules that exhibit distinct temporal activation patterns, with specific TF-gene circuits switching on and off as development progresses (Fig. 4h). These findings raise a fundamental question about transcription factor function: if regulatory modules activate and deactivate temporally, do individual transcription factors play different roles – or have different importance – at different developmental stages? To address this question systematically, we leveraged our time-resolved GRNs to perform computational genetic perturbations that could test how transcription factor knockouts affect cellular differentiation across the developmental timeline.

### *In silico* genetic perturbations suggest dynamic roles of transcription factors

With temporally dynamic regulatory modules, we next asked whether individual transcription factors exhibit stage-dependent functional roles. The regulatory module governing neuronal differentiation exemplifies this temporal specificity – being inactive at 10 hpf but activating by 14 hpf (Fig. 4c-bottom; see Interactive Web Portal), suggesting that factors like *pax6a* and *foxp4* acquire lineage-specifying functions as development progresses. To systematically test this hypothesis, we performed time-resolved *in silico* genetic perturbations across developmental stages.

Using CellOracle^27^, we simulated transcription factor knockouts by setting TF expression to zero and computing downstream effects on gene expression and cell fate trajectories (See Methods for details). We visualized perturbation effects through UMAP projections of cell-cell transition probabilities (Fig. a) and quantified them using cosine similarity between wild-type and perturbed states (Fig. b). This dual approach enabled both qualitative visualization and quantitative assessment of how TF knockouts alter the developmental trajectories of early neuro-mesodermal progenitors over time.

Systematic perturbation analysis revealed a striking developmental transition in TF function (Fig. b,c). Early in development (10 hpf), most transcription factors influenced both mesodermal and neuro-ectodermal lineages, reflecting high regulatory plasticity. However, by 24 hpf, TFs became largely lineage-committed, affecting either mesoderm or neuroectoderm but rarely both – a shift from broad regulatory influence to specialized lineage control.

The transcription factor *pax6a* exemplifies this functional transition. Known for its broad role in neural tube patterning and neuronal progenitor specification ^65,66^, *pax6a* showed dramatically different perturbation profiles across developmental time (Fig. d, e). At 10 hpf, *pax6a* knockout disrupts cell fate trajectories in both paraxial mesodermal and neuro-ectodermal lineages (Fig. d). By 19 hpf, however, predicted perturbation effects became restricted to the neuro-ectodermal lineage, with minimal impact on paraxial mesodermal development (Fig. e). This temporal refinement of function – from broad to specific – illustrates how transcription factors transition from having multi-lineage regulatory influence to enforcing lineage commitment.

These time-resolved *in silico* perturbations suggest that transcription factor function is inherently temporal: the same TF can act as a broad regulator early in development and a lineage-specific factor later, highlighting the importance of developmental context in defining gene function.

### A web resource for data exploration and timecourse *in silico* genetic perturbation

To facilitate access to our time-resolved single-cell multiome data and enable hypothesis generation by the broader research community, we developed Zebrahub-Multiome, an interactive web portal integrated into the existing Zebrahub resource^20^ (Fig. 6). This platform uniquely combines chromatin accessibility, gene expression, and regulatory network analyses in a unified interface, enabling researchers to seamlessly navigate between molecular modalities and developmental stages.

**Figure 6:**
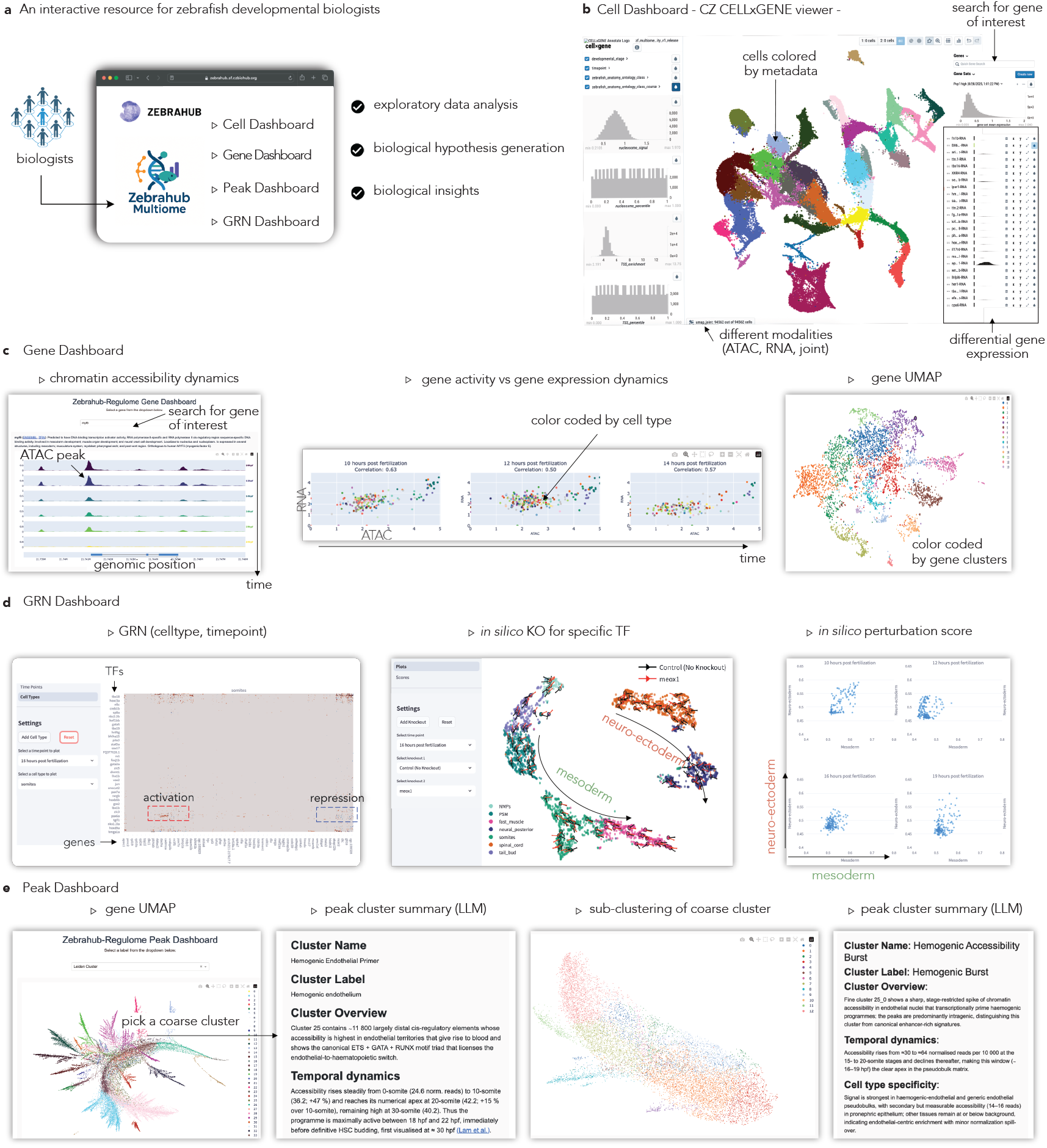
Zebrahub-Multiome: An interactive web resource for zebrafish developmental multi-omics exploration. **a**, Overview of the Zebrahub-Multiome web portal architecture, providing four specialized dashboards for comprehensive data exploration by the developmental biology community, enabling exploratory data analysis, biological hypothesis generation, and biological insights. **b**, Cell Dashboard powered by CZ CELLxGENE viewer enables interactive exploration of single-cell data with metadatabased cell coloring, selection of cell populations of interest, and differential gene expression analysis to identify tissue- and cell-type-specific marker genes. **c**, Gene Dashboard provides gene-centric analysis, including the chromatin accessibility coverage plot for the gene of interest, correlation analysis between gene activity and gene expression resolved by cell type and developmental stage, and gene-based UMAP visualization for exploring regulatory patterns. **d**, GRN Dashboard enables exploration of cell-type and stage-specific gene regulatory networks, with interactive *in silico* transcription factor knockout functionality and systematic quantification of perturbation effects across developmental trajectories. **e**, Peak Dashboard facilitates chromatin accessibility analysis through hierarchical peak clustering, allowing users to select coarse clusters, zoom into fine clusters, and access AI-powered peak cluster summaries using large language model (LLM) integration for automated functional annotation. The portal integrates all analyses presented in this study into an accessible interface, enabling researchers to explore zebrafish gene regulatory dynamics across developmental time and cell types without requiring computational expertise.

The portal supports multiple analytical workflows tailored for geneticists, molecular biologists, and developmental biologists (Fig. 6a). Users begin by exploring the cell atlas through the CZ CELLxGENE browser^67^, where they can visualize gene activity (ATAC) or expression (RNA) across tissues and developmental stages to identify cell-type-specific marker genes (Fig. 1b). These marker genes can then be analyzed in the Gene Dashboard, which tracks chromatin accessibility and expression dynamics across cell types and stages as demonstrated in Fig. 2. The dashboard also provides gene-level correlation analysis between chromatin accessibility and expression across the full transcriptome, and UMAP visualization for gene clusters with similar dynamic patterns across cell types and stages, enabling users to identify genes with similar regulatory patterns (Fig. 6c).

Beyond individual gene analysis, the portal enables exploration of gene regulatory networks at multiple scales. Users can investigate GRNs across developmental lineages or specific time points (Fig. 6d), with zoom capabilities for examining specific regulatory modules as presented in Fig. 4. Most uniquely, the platform supports *in silico* genetic perturbation experiments: after identifying transcription factors of interest through the Gene and GRN dashboards. For example, users can simulate their knockout and visualize predicted effects on cell fate trajectories for the paraxial mesoderm and anterior neuro-ectodermal lineages from neuro-mesodermal progenitors (Fig. 6e). This functionality enables the systematic comparison of perturbation effects across developmental stages and cell types, as illustrated in Fig..

Furthermore, the regulatory genomic elements, clustered by their accessibility over cell types and time points, are shared as a Peak Dashboard. Briefly, the peak UMAP visualization highlighting coarse and fine peak clusters with their LLM-derived annotation are shared so that the users can do a deeper dive into the dataset, opening a new door for biological hypothesis generation.

By integrating data visualization, regulatory network analysis, perturbation modeling, and characterization of regulatory elements in a single platform, Zebrahub-Multiome transforms our complex datasets into an accessible resource for generating and testing hypotheses.

## Discussion

We presented Zebrahub-Multiome, an integrated atlas of chromatin accessibility and gene expression during zebrafish embryonic development, capturing molecular states from 94,562 cells across six developmental stages spanning somitogenesis (10-24 hpf). By profiling both modalities from the same cells, we provide direct measurements of the relationship between regulatory element activity and transcriptional output during a critical period of vertebrate development.

Our analysis reveals that chromatin accessibility and gene expression follow distinct but coordinated temporal trajectories. Chromatin accessibility broadly decreases after 14-16 hpf while cell-type-specific gene expression increases, quantifying the progressive restriction of developmental potential observed across animal development^25,68,69^. This temporal offset between chromatin opening and gene activation, shown in Figure. 2f, has been documented in mammalian gastrulation, where accessibility changes precede expression by several hours^22^, and in human fetal development, where poised enhancers mark future cell fates^70,71^. Our measurements provide temporal resolution for this relationship across thousands of genes, revealing that genes associated with early somitogenesis show coordinated decreases while early organogenesis genes exhibit delayed activation following chromatin opening, consistent with the chromatin potential model^72^ (Fig. 2).

The analysis of 640,830 chromatin accessibility peaks revealed an unexpected hierarchical organization: regulatory elements group into modules based on their association with developmental pathways rather than genomic proximity. The finding that 77% of regulatory elements reside in intergenic and intronic regions with highly dynamic patterns while promoters remain constitutively accessible aligns with observations from mammalian development^49,52,53^ and suggests that promoters provide stable platforms while enhancers confer spatiotemporal specificity. The functional organization we observe – where peaks associated with specific signaling pathways cluster together regardless of chromosomal location – supports models of modular regulatory architecture that facilitate evolutionary innovation^73^ (Fig. 3).

The gene regulatory networks we infer show systematic changes during developmental progression. Network similarity steadily decreases, providing a quantifiable basis for the concept of progressive lineage restriction^1,2^. This quantitative measurement complements theoretical models of GRN evolution during development^74^. Early developmental stages share regulatory modules across multiple lineages – e.g. *hmga1a*-driven ribosomal biogenesis program at 10 hpf – that fragment into lineage-specific modules as development proceeds, consistent with the stratified organization of developmental GRNs^1^ (Fig. 4).

Our *in silico* perturbation experiments reveal that individual transcription factors can transition from broad, multi-lineage regulators to specialized, lineage-specific factors during development. The transcription factor *pax6a*, for instance, influences both mesodermal and neural trajectories at 10 hpf but becomes exclusively neural-specific by 19 hpf, exemplifying the concept of transcription factor redeployment in evolving cell types^75–77^. These computational predictions generate specific hypotheses about the temporal windows when transcription factors control cell fate decisions (Fig.).

By extracting subnetworks based on peak cluster organization, we reduced GRN complexity by 99% while preserving functionally coherent regulatory interactions (Fig. 4). This approach revealed temporally discrete regulatory circuits that would be obscured in whole-network analyses. The modular organization we identify may explain both developmental robustness and evolutionary flexibility, as modules can be gained, lost, or redeployed without disrupting entire regulatory architectures, a principle observed across metazoan evolution^73,74^ (Fig. 4).

Several limitations shape the interpretation of our findings. Sampling at discrete time points cannot capture continuous developmental processes or transient regulatory states. The computational inference of regulatory networks identifies statistical associations rather than direct regulatory interactions, and our predictions do not account for post-transcriptional regulation or non-cell-autonomous signaling. The loss of spatial information prevents us from determining whether regulatory programs overlap spatially. Future integration with spatial transcriptomics will be necessary to map regulatory states to tissue architecture^78,79^.

This resource complements existing zebrafish transcriptomic atlases^15–17,19,20^ and epigenomic atlases^38,46,80^ by jointly profiling gene expression and chromatin accessibility, and providing higher temporal resolution during somitogenesis. Notably, our study complements the recent work by Liu et al., ^81^, which characterized a single-cell multi-omic atlas of early zebrafish embryos from 3.3 to 12 hpf. Our data spans from 10 to 24 hpf, extending the temporal coverage of zebrafish embryonic development, benefiting the broader zebrafish research community. In the broader context of developmental biology, our study contributes to the growing body of work emphasizing the dynamic and context-dependent nature of gene regulation. Comparison with chromatin accessibility atlases from other vertebrates, including mouse^22,69^ and human^70,82,83^, will help distinguish conserved from species-specific regulatory strategies.

The web portal at zebrahub.org provides multiple entry points for data exploration, including gene- and peak-centric views, interactive cell embeddings, regulatory network visualization, and in silico perturbation predictions. The systematic annotation of regulatory elements using large language models transformed 77% of previously uncharacterized regions into functionally interpretable modules, providing candidates for targeted experimental perturbations.

In conclusion, Zebrahub-Multiome provides a quantitative atlas that captures the temporal coordination between chromatin accessibility and gene expression during zebrafish development. While our findings align with established principles – progressive restriction, modular organization, transcription factor redeployment – the value lies in providing quantitative resolution and systematically identifying the specific regulatory modules that implement these principles. By making these data and analyses freely accessible, we aim to accelerate both hypothesis-driven research and systematic exploration of vertebrate developmental regulation.

## Acknowledgements

We thank the entire Computational Biology & Genomics Platforms, and the Royer group of the Chan Zuckerberg Biohub-SF for helpful discussion and support. We thank the Scientific Computing team for their support for streamlining the data processing workflows. We appreciate the brainstorming and feedback from Luca Pinello, Max Lombardo, Sara Simmonds, Alec Tarashansky, Pablo Garcia-Nieto, Guillaume Le Treut, Andy Zhou, Bohyeon Yu, Liliana Solnica-Krezel, Will Greenleaf, Mathias Voges, Manuel Lionetti, and Kenji Kamimoto. We thank Sandra Schmid, Scott Fraser, Joseph DeRisi, Bill Burkholder, Giovanni Palla, Danny Conrad, Yuening Liu, and Vera Janssen for the critical reading and feedback on the manuscript. Funding for this work was provided by Chan Zuckerberg Biohub San Francisco (CZB SF). We thank the CZB SF donors, Priscilla Chan and Mark Zuckerberg, for their generous support.

## Data availability

Raw sequencing data are available at NCBI’s SRA BioProject (PRJNA1164307). The processed h5ad objects as well as Oracle objects (for GRNs) are available in Google Drive. The dataset can be interactively explored at zebrahub.org.

## Code availability

The code/notebooks used for data processing and analysis are in github repository: https://github.com/czbiohub-sf/zebrahub-multiome-analysis

## Methods

### Zebrafish husbandry

The zebrafish care and experimental protocols adhered to guidelines approved by the institutional animal care and use committee (IACUC) at the University of California San Francisco (UCSF). Wild-type EKW embryos were utilized for the multiome experiments. Adults zebrafish were bred and maintained at a temperature of 28.5°C, receiving food three times daily through an automatic feeder. Embryos were raised and staged according to the somite count, along with the hours post-fertilization (hpf), serving as temporal phenotypic markers during somitogenesis.

### Single-cell multiome sample preparation

For multiome experiments, 60 embryos were pooled for each developmental stage. We have one biological replicate for 15 somites (16 hpf) as an internal control. First, the embryos were dechorionated using pronase solution (1mg/ml). The embryos were incubated in pronase solution for a maximum of 2 minutes. This is followed by a series of washes using egg water till the chorions are removed and any remnant pronase solution is washed off. Next, 60 embryos at the required stage were dissociated, and their nuclei were isolated using the 10x Nuclei Isolation Kit (PN-1000494). The nuclei were resuspended in 25 ul of resuspension buffer. The nuclei count was determined using a dilution (1:4) of the cell suspension and DAPI solution (1:1000 dilution). The cells are incubated in DAPI for 15 minutes at room temperature in the dark. This is followed by counting nuclei manually using a hemocytometer.

### Single-cell multiome library preparation and sequencing

The single-nuclei suspensions were transposased such that the transposase enters the nuclei and preferentially fragments the DNA in open regions of the chromatin. This is then processed into droplet emulsions using the Chromium Single Cell Controller (10x Genomics). Both scRNA-seq libraries and ATAC libraries are generated for each cell using the Chromium Next GEM Single Cell Multiome ATAC + Gene Expression reagents. The single-nuclei suspensions were assessed using the hemocytometer to determine the nuclei count, we aimed for a maximum recovery of 10,000 nuclei wherever possible. Fragmentation, ligation, and PCR amplification were performed using the BioRad C1000 Touch Thermal Cycler with thermal profiles specified by the 10X Genomics User Guide CG000338. Amplified cDNA and final ATAC, gene expression libraries were assessed using the Agilent 4200 TapeStation/High Sensitivity D5000 Screentape and qPCR with the Kapa Library Quantification kit for Illumina. Each library was normalized to 4 nM and pooled for each NovaSeq sequencing run. The pools were sequenced using Novaseq run kits, with either single or dual index plates (Illumina 20028313), and a 1% PhiX control library was spiked in. Sequencing was performed on the NovaSeq 6000 Sequencing System (Illumina) aiming for 1 billion reads per library.

### Single-cell Multiome data processing

Sequencing reads were aligned to the Ensembl reference genome (GRCz11) using Cellranger-arc (version 2.0.2, 10x Genomics), generating count matrices for both RNA (Gene Expression) and ATAC modalities. Subsequent processing of these raw count matrices was performed using the Seurat (version 4.4.0) and Signac (version 1.11.0) packages^37,41^. For each developmental stage, we created Seurat objects incorporating both “RNA” and “ATAC” assays. Low-quality nuclei were filtered out based on the following quality control (QC) metrics:

- RNA modality: Cells were retained if they contained between 1,000 and 25,000 UMIs.
- ATAC modality: Cells were retained if they contained between 1,000 and 100,000 fragments, exhibited a nucleosome signal below 2, and demonstrated a TSS enrichment signal above 1.

The RNA modality was processed first by normalizing the raw transcript counts using counts per million (CPM), followed by scaling and principal component analysis (PCA). This process resulted in the “SCT” assay, generated using the *scTransform* function from the Seurat package.

### Peak calling in scATAC-seq datasets

In scATAC-seq analysis, the features of interest are “peaks,” which represent discrete regions of open chromatin in the genome, in contrast to the “genes” analyzed in scRNA-seq. Peak calling in scATAC-seq data presents unique challenges, particularly in determining how to group cells with similar chromatin accessibility profiles to achieve robust peak identification^41^. We employed MACS2, as implemented in Signac, to call peaks using two different grouping strategies: (1) “cell-type” grouping, based on annotations transferred from Zebrahub, and (2) “bulk” grouping, considering all cells collectively. This approach, combined with the peaks called by Cellranger-arc, resulted in three distinct peak profiles (Figure S4a). To integrate these profiles, we employed the “iterative peak calling” protocol from ArchR, which prioritizes peaks based on higher confidence. To create a comprehensive atlas of cell-type-specific regulatory elements, we implemented the following strategy: We first established celltype-specific peaks as the baseline. We then incorporated nonoverlapping peaks from the bulk analysis. Finally, we iteratively added Cellranger-arc peaks that were absent from the previous two profiles (Fig. S4b).

### Iterative peak calling strategy

Our analysis revealed that MACS2-called peaks were generally shorter than those identified by Cellranger-arc, suggesting superior detection sensitivity for MACS2 (Fig. S4c)^41^. The number of peaks added at each iteration (Fig. S4d) indicated substantial overlap between cell type-specific and bulk peaks called by MACS2 with those detected by Cellranger-arc. This iterative peak-calling approach offers two key advantages: it captures peaks specific to rare cell types that may be missed in bulk-level analyses, and it identifies specific regulatory regions that enhance the identification of transcription factor binding motifs in subsequent analyses.

### Gene activity score quantification

Gene activity scores were computed using Signac, which sums up the scATAC-seq reads within the gene body and 2kb upstream of the transcription start site for each gene^41^. The resulting gene activity scores were log-normalized to account for the sequencing depth for downstream analyses.

### Individual dataset dimensionality reduction and embedding

Following peak calling and gene activity score computation, individual datasets underwent dimensionality reduction for both modalities. For RNA, principal component analysis (PCA) was performed on the SCT-normalized data using the top 50 components. For ATAC, the merged peak set (`peaks merged` assay) was processed using term frequency-inverse document frequency (TF-IDF) normalization, followed by singular value decomposition (SVD) to generate latent semantic indexing (LSI) embeddings, excluding the first component to avoid sequencing depth artifacts^68,69^. Individual dataset embeddings were computed using UMAP on PCA (RNA, dimensions 1-50) and LSI (ATAC, dimensions 2-40) representations. Weighted nearest neighbor (WNN) graphs were constructed using the FindMultiModalNeighbors function in Seurat to integrate RNA and ATAC modalities at the single-dataset level, generating joint UMAP embeddings that capture both transcriptomic and epigenomic similarity within each timepoint ^37^.

### Cross-dataset integration and batch correction

To integrate data across the six developmental time points while correcting for batch effects, we implemented separate integration workflows for RNA and ATAC modalities. First, a unified peak set was created across all datasets using the UnifyPeaks function with mode=“reduce”, generating a consensus set of accessible chromatin regions. Count matrices were recomputed for each dataset using the unified peak set via the FeatureMatrix function, ensuring consistent feature spaces across time points. For ATAC integration, we employed reciprocal LSI (rLSI) as implemented in Signac’s FindIntegrationAnchors function. Integration anchors were identified using all unified peaks as anchor features, with LSI dimensions ranging from 2 to 30 used for anchor identification. The IntegrateEmbeddings function was then applied to generate batch-corrected LSI embeddings (`integrated lsi`), enabling cross-dataset comparison while preserving biological variation. For RNA integration, we used reciprocal PCA (rPCA) with SelectIntegrationFeatures to identify highly variable genes across datasets. Integration anchors were computed using FindIntegrationAnchors with normalization.method=“LogNormalize”, reduction=“rpca”, and reference time points (10 hpf and 24 hpf) to guide the integration process. The IntegrateData function generated batchcorrected expression values (“integrated” assay), which were scaled and subjected to PCA to produce integrated RNA embeddings (`integrated lsi`).

### Joint multimodal embedding computation

Integrated RNA and ATAC modalities were combined using weighted nearest neighbor (WNN) analysis to generate unified cell embeddings that leverage both transcriptomic and epigenomic information. The FindMultiModalNeighbors function from Seurat^37^ was applied using the `integrated pca` (dimensions 1-30) and `integrated lsi` (dimensions 2-40) embeddings as input, with modality-specific weights computed based on the relative contribution of each modality to local neighborhood structure. The resulting WNN graph was used to compute a joint UMAP embedding (“wnn.umap”) that reflects both gene expression and chromatin accessibility patterns across all developmental time points. Cell-specific modality weights were stored as metadata to enable assessment of whether individual cells were better characterized by their transcriptomic or epigenomic profiles. Leiden clustering was performed on the WNN graph to identify transcriptionally and epigenomically coherent cell populations for cell type annotation.

### Cell type annotation

Cell type annotation was performed on the joint UMAP embedding using a multi-step workflow that leverages both computational label transfer and expert biological curation. Initial cell type predictions were obtained using scANVI^84^ to transfer labels from the Zebrahub reference transcriptomic atlas^20^ to our dataset using the RNA modality. Additional reference atlases including Daniocell^85^ and Zscape^18^ were consulted for validation purposes.

To refine these initial predictions, we performed Leiden clustering on the weighted nearest neighbor (WNN) graph, which captures cellular similarities based on both modalities^37^. Zebrafish biologists then performed expert curation by analyzing differential gene expression patterns between Leiden clusters using scanpy’s t-test implemented in CZ CELLxGENE Annotate ^67^). This differential expression testing, combined with established RNA marker genes^20^ and zebrafish ontology terms from ZFIN, guided decisions to merge over-segmented clusters or subdivide heterogeneous clusters to achieve biologically meaningful cell type annotations at both coarse and fine resolution levels, as illustrated in Fig. 1b-d. This iterative workflow ensures that final annotations capture the discrete cellular identities revealed through multimodal integration while being grounded in established zebrafish developmental biology knowledge.

### Tissue-specific marker gene visualization

To visualize and validate the cell type annotations in our single-cell atlas, we selected sets of well-established marker genes for four major tissue types: endoderm, mesoderm, neuroectoderm, and hematopoietic lineages. The expression patterns of these markers were overlaid on the UMAP embeddings to confirm the identity of annotated cell clusters and to illustrate the spatial organization of different cell types in the low-dimensional space. For endodermal tissues, we used the following marker genes: *fgfrl1b, col2a1a, ptprfa, emid1, nr5a2, ism2a, pawr, mmp15b, foxa3*, and *onecut1*. Mesodermal markers included *msgn1, meox1, tbx6, tbxta, fgf8a*, and *her1*. To identify neuroectodermal tissues, we employed *pax7a, pax6a, pax6b, col18a1a, en2b, znf536, gpm6aa, gli3*, and *chl1a* as marker genes. For each tissue type, we calculated the average log-normalized expression of its respective marker genes for each cell. These average expression values were then projected onto the UMAP embedding, with color intensity representing the level of expression of the marker gene. This approach allowed us to visualize the spatial distribution of different tissue types within the UMAP space and validate our cell type annotations.

### Cell type coarse-graining into developmental lineages

To facilitate lineage-level analysis, we grouped the fine-grained cell type annotations into broader developmental lineages based on their embryological origins and developmental trajectories. The mapping from individual cell types to lineages is summarized in Table 1.

### Neighborhood graph comparison between different modalities

Cross-modality validation was performed to assess how well each modality preserves biological structure using two complementary approaches: neighborhood purity analysis and trajectory conservation analysis.

First, neighborhood purity analysis was conducted using the connectivity matrices for RNA (*RNA*_*con*_), ATAC (*ATAC*_*con*_), and joint (*con*_*wnn*_) neighborhood graphs. For each cell, neighborhood purity was calculated as the proportion of *k*-nearest neighbors (*k* = 30) sharing the same metadata label (either cell type annotation or Leiden clusters computed from each modality). Purity scores were computed for three clustering approaches: RNA Leiden clusters (resolution=0.8), ATAC Leiden clusters (resolution=0.5), and joint Leiden clusters (resolution=0.35), resulting in roughly 23-27 Leiden clusters for each modality. Results were visualized using box plots to reveal the differences in neighborhood consistency across modalities.

Second, trajectory conservation analysis quantified preservation of developmental ordering along established NMP-derived trajectories: mesodermal (NMPs → tail bud → PSM → somites → fast muscle) and neuro-ectodermal (NMPs → spinal cord → neural posterior). For each modality, diffusion pseudotime was computed using scanpy’s diffusion map algorithm ^86^ with NMPs as root cells. Trajectory conservation scores were calculated by measuring Pearson correlation between observed pseudotime ordering and expected developmental sequence to the −1 to +1 range, where 1 indicates perfect preservation of developmental ordering, and −1 indicates incorrect ordering. To ensure numerical stability, analyses were performed on subsampled datasets (*n* = 10000 cells) with matrix regularization to handle disconnected graph components.

### Metacell inference

To account for the noise and sparsity of single-cell Multiome datasets for the correlative analysis of chromatin accessibility and gene expression, we used SEACells^42^ to compute the metacells. Specifically, for each dataset, we utilized the ATAC modality to compute latent semantic indexing (LSI) as the dimensionality reduction method. We used the 2nd to 40th LSI components to compute the metacells, and we excluded the first LSI component as the it is usually correlated with the sequencing depth. After the metacell computation, we aggregated the gene expression (RNA) and gene activity (ATAC) values at the metacell level, creating a set of aggregated count matrices used for Fig. 2. The number of cells per metacell was set to 75 following the SEACells tutorial^42^. Note that we used 30 cells/metacell for the Fig. analysis for a fine-grained visualization of the cell-cell transition probability in 2D embeddings.

### Gene dynamics matrix construction

To systematically analyze temporal patterns of gene regulation across development, we created gene dynamics matrices that capture both chromatin accessibility and changes in gene expression across cell types and developmental stages. First, metacells were computed using SEACells^42^ to aggregate transcriptionally and epigenomically similar cells, thereby reducing noise while preserving cellular heterogeneity (see the Metacell Inference section). For each developmental time point, we computed the mean gene expression (RNA) and mean gene activity scores (ATAC) across metacells belonging to the same annotated cell type. This resulted in two matrices per timepoint: an RNA expression matrix and an ATAC accessibility matrix, each with genes as rows and annotated cell types as columns. To create the final gene dynamics matrix, we concatenated the mean expression and accessibility values across all six developmental time points (10, 12, 14, 16, 19, and 24 hpf) for each gene. Each gene was thus represented by a high-dimensional vector capturing its regulatory dynamics across the entire developmental landscape, with dimensions corresponding to:

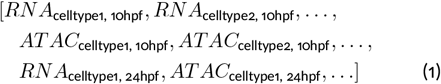

This approach enabled the comparison of genes based on their complete temporal and cell-type-specific regulatory profiles.

### Gene UMAP computation

The gene dynamics matrix was filtered to retain highly variable genes for either RNA or ATAC modality (n=5,068) and subjected to dimensionality reduction using Principal Component Analysis (PCA), retaining the top principal components that explained the majority of variance. Uniform Manifold Approximation and Projection (UMAP) was then applied to the PCA embeddings (using 50 principal components) to generate a two-dimensional representation, where each point represents a gene positioned according to its regulatory dynamics similarity. Leiden clustering was performed on the gene UMAP to identify functionally related gene modules. Clustering resolution was optimized to balance cluster granularity with biological interpretability. The resulting gene clusters were analyzed for functional enrichment using FishEnrichr^43,44^ to identify associated developmental pathways and biological processes. This gene-centric approach enabled the discovery of coordinated regulatory programs that would not be apparent from traditional expression-based clustering approaches.

### Gene dynamics pattern analysis

To quantify the temporal behavior of individual genes across development, we performed a systematic analysis of temporal patterns in the gene dynamics matrices. For each gene, we first aggregated the mean expression (RNA) and mean chromatin accessibility (ATAC) values across all cell types for each developmental timepoint, creating embryo-wide temporal profiles that capture the overall regulatory trajectory of each gene throughout the 10-24 hpf developmental window. We then performed linear regression to quantify the temporal trends in gene expression and chromatin accessibility. For each gene, separate linear regression models were fitted to the aggregated RNA and ATAC values across the six developmental time points (10, 12, 14, 16, 19, and 24 hpf). The slope coefficients from these models provided quantitative measures of temporal change, with positive slopes indicating increasing activity over time, negative slopes indicating decreasing activity, and slopes near zero indicating either stable expression patterns or genes with peaks/troughs at intermediate time points. These slope values were used to classify genes into temporal pattern categories and were visualized on the gene UMAP to reveal the spatial organization of genes with similar temporal behaviors. To distinguish between truly stable genes and those with intermediate peaks or troughs (both of which can exhibit slopes near zero), we determined the developmental stage at which each gene shows maximum activity. For each gene, we identified the time point of peak expression and peak chromatin accessibility by finding the time point with the highest aggregated value across the developmental window. This peak timing information was used to color-code genes on the UMAP visualization, revealing temporal waves of gene activation during embryonic development and enabling discrimination between stable expression patterns and transient activation patterns. To ensure that visualized patterns represent significant biological signals rather than noise, we computed a peak contrast metric for each gene, defined as follows: (peak value - mean of other time points) /standard deviation of all time points, excluding the maximum point. This peak contrast value was used as the transparency (alpha) for each gene in the UMAP visualization, with genes showing stronger temporal patterns (higher contrast) displayed with greater opacity and genes with weaker or potentially noisy patterns shown with increased transparency.

This approach emphasized genes with robust temporal dynamics while de-emphasizing those with minimal or inconsistent temporal changes.

### Gene set enrichment analysis

To identify the biological pathways enriched in a group of genes identified from clustering, we used FishEnrichr, a web-based gene set enrichment analysis tool, specialized for zebrafish^43,44^. We used the list of genes identified from the clustering and referred to the gene ontology terms of enriched pathways from WikiPathways database^87^.

### Comprehensive pseudobulk chromatin accessibility analysis across cell types and time points

To assess the dynamics and variability of chromatin accessibility patterns across different cell types and developmental time points, we performed pseudobulk aggregation of all ATAC-seq peaks from the single-cell data.

Single-cell accessibility profiles were aggregated (“pseudobulked”) by grouping cells according to their Leiden clusters from the joint UMAP and developmental time points. For each combination of cell cluster *i* and time point *j*, all cells belonging to that (*i, j*) group were pooled, and the read counts (accessibility signals) per peak were summed. This generated one “pseudobulk” accessibility vector per cell cluster-timepoint combination, effectively reducing sampling variability and highlighting population-level differences.

Each pseudobulk profile was normalized by a scaling factor (total read counts per group divided by the median of total read counts across all groups) to account for sequencing depth and differences in cell numbers between groups. We then transposed the normalized count matrix to create a peaks-by-pseudobulk matrix, where each row represented a genomic peak and each column represented a cell cluster at a specific time point.

Principal Component Analysis (PCA) was applied to project all peaks into a lower-dimensional space, retaining the top principal components that explained the majority of variance. Uniform Manifold Approximation and Projection (UMAP) was performed on these PCA embeddings (using 1-40 principal components, parameters *n neighbors* = 15 and *min dist* = 0.3) to obtain a two-dimensional layout. Each point in the UMAP corresponds to a single genomic peak, and clustering reflects similarities in accessibility patterns across cell clusters and developmental stages.

### Annotation of peak types using genome annotation

Peak genomic annotations were assigned using the Ensembl zebrafish genome annotation (GRCz11). Each peak was classified into one of four categories based on its genomic location: promoter peaks (within 500bp upstream or 100bp downstream of transcription start sites), exonic peaks (overlapping with annotated exons), intronic peaks (located within annotated introns), and intergenic peaks (located outside of annotated gene boundaries) ^23^. The distance to the nearest transcription start site (TSS) was calculated for all peaks, with a maximum of 50 kbp, and the distribution of these distances was analyzed to understand the spatial organization of regulatory elements.

### Annotation of peaks with associated genes

To link regulatory peaks to their target genes, we employed a two-step annotation approach that prioritizes functional relationships over genomic proximity. First, we computed correlation-based peak-gene associations by calculating Pearson correlation coefficients between peak accessibility (ATAC-seq) and gene expression (RNA-seq) across all cells using Signac^41^. Peak-gene pairs with correlation coefficients r *>* 0.3 and FDR *<* 0.05 were considered functionally linked, regardless of genomic distance. Second, for peaks not assigned through correlation analysis, we used a genomic overlap-based annotation method, linking peaks to genes whose gene bodies (including exons and introns) overlapped with the peak coordinates. Peaks within 2kb upstream of transcription start sites were also included in this genomic proximity-based assignment.

When conflicts arose between correlation-based and overlap-based annotations (i.e., a peak showed significant correlation with one gene but overlapped with a different gene), the correlation-based assignment took priority, as it reflects functional regulatory relationships rather than simple genomic proximity. This hierarchical approach ensured that peaks were associated with genes based on evidence of regulatory activity while providing genomic fallback assignments for peaks without detectable correlation signals.

### Hierarchical clustering of peak UMAP

To identify functionally related groups of regulatory elements, we performed hierarchical clustering on the peak UMAP embeddings using the Leiden algorithm for clustering. Multiple clustering resolutions were tested to achieve an optimal balance between cluster granularity and biological interpretability. The final clustering resulted in 36 major peak clusters. For fine-grained analysis, sub-clustering was performed within major clusters showing high internal heterogeneity, generating additional sub-clusters for detailed functional characterization. Clusters with fewer than 100 peaks were merged into the nearest cluster using the neighborhood graphs.

To annotate peak clusters with their dominant celltype identity, we developed a contrast-weighted enrichment analysis that prioritizes celltype-specific regulatory elements over broadly accessible peaks. For each peak, we calculated celltype contrast as a z-score: (*max*_*acc*_ − *mean*_*others*_)*/std*_*others*_, quantifying how specifically accessible each peak is for its assigned celltype.

### Differential motif enrichment analysis

To identify transcription factor (TF) motifs differentially enriched within chromatin accessibility clusters, we employed *GimmeMotifs*’ Maelstrom module, a comprehensive computational framework for motif activity analysis ^88,89^. Peaks were first grouped using Leiden clustering based on their chromatin accessibility profiles across cell types and developmental stages, with a resolution of 0.7. Motif scanning was performed on all peak sequences extracted from *danRer11* reference genome using position weight matrices (PWMs) from the CisBP motif database (curated for zebrafish, termed `CisBP danio rerio v2`) ^27^. The resulting motif scores were used for differential enrichment analysis across clusters in Maelstrom. Motif activity per cluster was assessed through multiple statistical and machine-learning approaches, including Mann–Whitney U tests, hypergeometric enrichment, random forest classification, and Bayesian ridge regression. To reduce redundancy in the results, highly correlated motifs (correlation *>* 0.8) were grouped, with only the most informative representative motif retained per group. Final motif significance was determined through rank aggregation of these methods, retaining only motifs that met stringent statistical criteria (*p <* 0.05, |*z*| *>* 2, and occurrence in at least 1% of peaks). The resulting motif-by-cluster z-score matrix revealed distinctive patterns of TF motif enrichment across accessibility clusters, providing insights into the regulatory factors that may drive cell-type-specific and temporally dynamic chromatin states during development.

### Annotation of peak clusters using Large language models

We developed a systematic annotation pipeline using Large Language Models (LLMs) to functionally characterize peak clusters identified through Leiden community detection. For each chromatin accessibility cluster, we compiled comprehensive feature sets including: (1) cluster-specific statistics (number of peaks, genomic annotations, median distance to TSS), (2) pseudobulk accessibility profiles across cell-type × time-point combinations with mean and standard error values, (3) number of cells per pseudobulk group for normalization context, (4) associated genes determined by correlation analysis or genomic overlap, (5) enriched transcription factor binding motifs with z-scores from MAELSTROM analysis, and (6) corresponding transcription factors from the CisBP database with consensus sequences and experimental validation status. These multi-modal datasets were formatted into structured queries and processed using Litemind (v2025.7.1) with OpenAI and Anthropic models, GPT-o3-high and Sonnet-4, respectively, equipped with web search capabilities and access to biological databases (Alliance, Ensembl, PubMed). The LLM agents were prompted to analyze temporal dynamics, cell-type specificity, and regulatory programs following standardized templates, with outputs subject to automated review and revision cycles. To ensure annotation quality and consistency, we employed a multi-tier approach with independent queries for each cluster, followed by systematic validation against external literature sources. This LLM-based pipeline enabled comprehensive functional characterization of hundreds of regulatory clusters while maintaining objectivity and reproducibility in the annotation process.

### Subsetting for the Neuro-Mesodermal Progenitor population

To investigate key transcription factors in Neuro-Mesodermal Progenitors (NMPs), we created a subset of our data focusing on NMPs and their subsequent developmental trajectories. This subset included cells from mesodermal (tail bud, presomitic mesoderm (PSM), somites, and fast muscle) and neuroectodermal (spinal cord, neural posterior) lineages.

### Pseudotemporal trajectory analysis for NMP-derived lineages

To analyze the temporal progression of gene expression along developmental trajectories, we performed pseudotemporal ordering of cells from neuro-mesodermal progenitor (NMP) populations and their derivatives.

Pseudotime inference was performed using Slingshot^90^, implemented via the Python package pySlingshot (https://github.com/mossjacob/pyslingshot). We used the 2D UMAP embedding as input for trajectory inference to compute the principal curve. NMPs were designated as the root cell type^20^, representing the starting point of both lineage trajectories. Slingshot was configured to identify bifurcating trajectories emanating from the NMP root, capturing the divergence into neuro-ectodermal and mesodermal lineages.

For each identified trajectory, pseudotime values were assigned to cells based on their position along the inferred developmental path. NMPs received pseudotime values near zero, and terminally differentiated cells (fast muscle for the mesodermal trajectory and neural posterior for the neuroectodermal trajectory) received maximum pseudotime values. Gene expression patterns were then visualized along these pseudotemporal axes by computing moving averages of normalized gene expression values within sliding windows of pseudotime, enabling identification of genes with stage-specific activation patterns during lineage specification (Fig. 1e).

### Chromatin co-accessibility computation

We used Cicero^91^ to calculate chromatin co-accessibilities for all peak-peak pairs within each chromosome. The resulting dataframe contained peak pairs (*i* and *j*) and their co-accessibility scores. We applied CellOracle’s filtering criteria:

a. Removed peak-peak pairs with co-accessibility scores below 0.8.
b. Retained pairs where at least one peak corresponded to a transcription start site (TSS, as annotated by Ensembl).

### Gene Regulatory Networks inference

We used CellOracle to infer cell-type specific Gene Regulatory Networks (GRNs) for the NMP-subsetted objects at each time point^27^. The process followed the CellOracle protocol as illustrated in Fig. S16. Briefly, CellOracle computes the distal and proximal *cis*-regulatory regions for each gene using cicero^91^. Then, transcription factor binding motifs were scanned within these *cis*-regulatory regions, constructing a base GRN, where the rows are all the activities of transcription factors, and binarized elements for the presence or absence of TF binding motif within the *cis*-regulatory element for that gene. Importantly, these base GRNs were computed for each time point, noting that the *cis*-regulatory elements change dynamically over developmental stages (Fig. S15). Using the base GRN, CellOracle constructed linear equations modeling each gene’s expression level as a combination of regulatory transcription factors from the base GRN. Coefficients for these models were determined through regularized linear regression, using gene expression values to approximate transcription factor activity. The resulting cell-type specific GRNs were further refined by filtering out weak edges based on edge strength and/or regression p-values (Fig. S16). Of note, we made a quantitative comparison of GRNs inferred from two biological replicates at 16 hpf. The network metrics were very similar between the biological replicates and the GRNs inferred from other time points (Fig. S18). Therefore, we used one of the replicates (TDR118) as a representative GRN for the downstream analyses.

### Quantification of GRN similarity across developmental stages and cell types

To quantify GRN evolution during development, we computed cosine similarity between all pairwise combinations of cell-type-specific GRNs using edge strength vectors. For cross-cell-type analysis, we calculated similarity between all 903 unique cell type pairs at each developmental stage. For temporal analysis, we computed similarity between GRNs from the same cell type across all timepoint pairs. Cell types were grouped into seven major lineages according to Table 1 for lineage-specific analysis. scikit-learn(python) was used to compute the cosine similarity^92^.

### Computing subGRN for each peak cluster

Peak clusters were analyzed for transcription factor motif enrichment using GimmeMotifs Maelstrom on fine-resolution Leiden clusters. Motif enrichment z-scores were thresholded at ≥2.0 to identify significantly enriched motifs per cluster. Threshold sensitivity analysis was performed across multiple values (0.5–3.0) to optimize signal-to-noise ratio while maintaining biological interpretability. Motifs were mapped to transcription factors using the CisBP v2.0 database. We developed a four-step computational pipeline to create regulatory program “meshes”—binary matrices representing predicted TF-gene regulatory relationships per peak cluster:

1. **Motif extraction**: Enriched motifs per cluster were identified using z-score thresholding (≥2.0)
2. **TF mapping**: Motifs were mapped to transcription factors via CisBP v2.0 annotations
3. **Gene association**: Linked genes were extracted per cluster using Signac correlation-based peak-to-gene linkages rather than simple genomic overlap to ensure regulatory relevance
4. **Matrix construction**: Binary TF-gene matrices were generated where matrix element (*i, j*) = 1 indicates TF *i* is predicted to regulate gene *j* within the regulatory program

Sub-GRNs were extracted by intersecting regulatory program meshes with cell-type and timepoint-specific gene regulatory networks from CellOracle. For each peak cluster, predicted TF-gene pairs from the mesh were filtered against the corresponding GRN using the following criteria: (1) presence in the regulatory program mesh and (2) presence in the CellOracle GRNs. This approach yielded context-specific sub-GRNs, representing the subset of predicted regulatory relationships that were actually implemented in each cellular context.

Cluster distinctiveness was assessed using Jaccard similarity analysis between peak clusters. Threshold optimization was performed using multiple statistical measures including mean/median motifs per cluster, proportion of clusters losing signal, and distribution analysis via histograms with kernel density estimation. Correlation analysis between TF-based and gene-based cluster similarities validated the biological coherence of the regulatory program definitions.

To visualize the dynamics of network, we generated a master layouts that encompassess all nodes and edges from sub-GRNs across all time points. This approach enabled systematic identification and temporal tracking of regulatory programs from chromatin accessibility data, providing insights into the dynamic implementation of gene regulatory networks during zebrafish embryonic development.

### Aligned UMAPs of individual time points

To visualize the temporal progression of cellular trajectories in NMP trajectories across time points, we used AlignUMAP to align the UMAPs from individual time points using pairwise anchors between time points. Briefly, the pairwise anchors were computed by nearest neighbors using the dimensionality reduction from the ATAC modality (Latent Semantic Indexing, LSI, from 2nd to 40th components). To avoid over-alignment, we relaxed the strength of the alignment regularization parameter to 0.001. The output is a series of UMAPs for each time point, which shows conserved orientation and configurations for easier visual inspection of the WT and *in silico* KO transitions.

### *in silico* genetic perturbation

We performed *in silico* genetic perturbation following CellOracle’s documentation. Briefly, CellOracle utilizes cell-type-specific gene regulatory networks and developmental trajectories to predict shifts in cellular differentiation vectors. First, it simulates the transcriptomic change upon knockout of the gene of interest by using signal propagation. Then, the changed transcriptome is compared with the transcriptome of cells in the local neighborhood to estimate the cell-cell transition probabilities.

### cell-cell transition probability inference for wild-type embryos

Notably, we used pySlingshot, a python implementation of Slingshot^90^) to compute the pseudotime with “NMPs” defined as the root cells^20^. Next, we used CellRank2^93^‘s “PseudotimeKernel” to compute the cell-cell transition probability matrix and 2D projections onto the embeddings (UMAP) for the wild-type (WT) trajectory inference.

## Supplementary Information

3) Figure for GRN comparison - biological replicates

**Figure S1:**
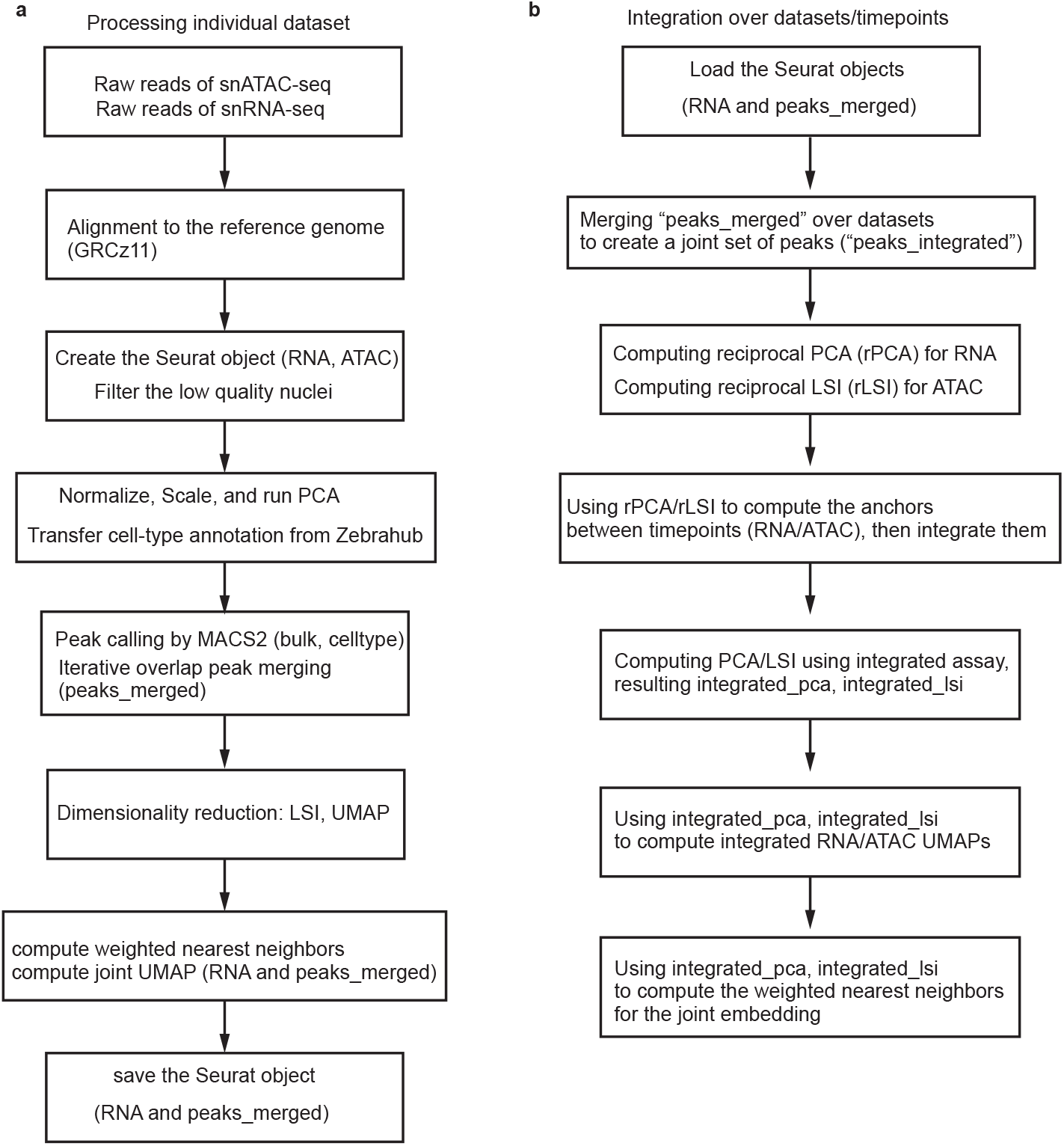
Workflow for processing and integrating snATAC-seq and snRNA-seq data Workflow for processing and integrating snATAC-seq and snRNA-seq data. **a**, Processing individual datasets: The workflow begins with raw reads from snATAC-seq and snRNA-seq, which are aligned to the zebrafish genome (DanRer11). Seurat objects are then created for both RNA and ATAC data, with low-quality nuclei filtered out. The data undergoes normalization, scaling, and PCA, followed by cell-type annotation transfer from Zebrahub. Peak calling is performed using MACS2, with subsequent iterative overlap peak merging. Dimensionality reduction techniques, including LSI and UMAP, are applied. The process concludes with the computation of weighted nearest neighbors and a joint UMAP for RNA and merged peaks, resulting in a final Seurat object that contains both RNA and merged peak data. **b**, Integration over datasets/time points: This phase begins by loading the previously saved Seurat objects. A joint set of peaks (“peaks integrated”) is created by merging across datasets. The workflow then employs reciprocal PCA (rPCA) for RNA and reciprocal LSI (rLSI) for ATAC data. These are used to compute anchors between time points and integrate the data. Following integration, PCA/LSI is computed using the integrated assay. The process culminates in generating integrated RNA/ATAC UMAPs using integrated PCA and LSI, and computing weighted nearest neighbors for the joint embedding.

**Figure S2:**
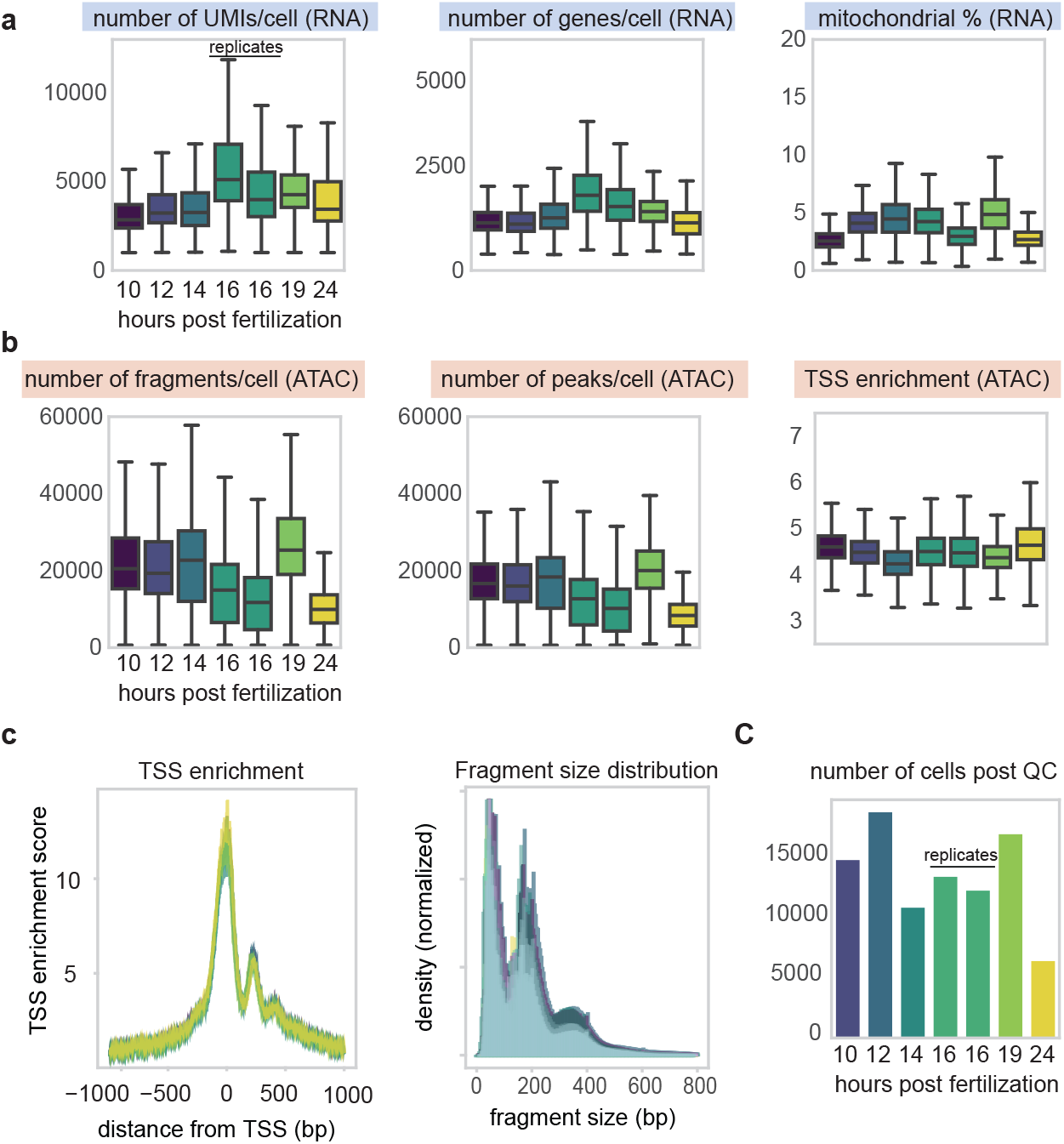
Quality control metrics for scRNA-seq and scATAC-seq datasets across zebrafish developmental time points. **a**, scRNA-seq metrics: Box plots show the distribution of (left) number of UMIs per cell, (middle) number of genes detected per cell, and (right) percentage of mitochondrial reads across developmental time points from 10 to 24 hours post fertilization (hpf). These metrics indicate the quality and depth of RNA sequencing data. **b**, scATAC-seq metrics: Box plots display (left) the number of fragments per cell, (middle) the number of peaks detected per cell, and (right) the transcription start site (TSS) enrichment score across the same developmental time points. These metrics reflect the quality and information content of ATAC sequencing data. **c**, Additional ATAC-seq quality metrics: (Left) TSS enrichment profile showing the aggregated ATAC-seq signal around transcription start sites. (Middle) Fragment size distribution of ATAC-seq reads, typically showing nucleosome-free regions and mono-nucleosome peaks. (Right) Bar plot of the number of cells passing quality control for each developmental timepoint.

**Figure S3:**
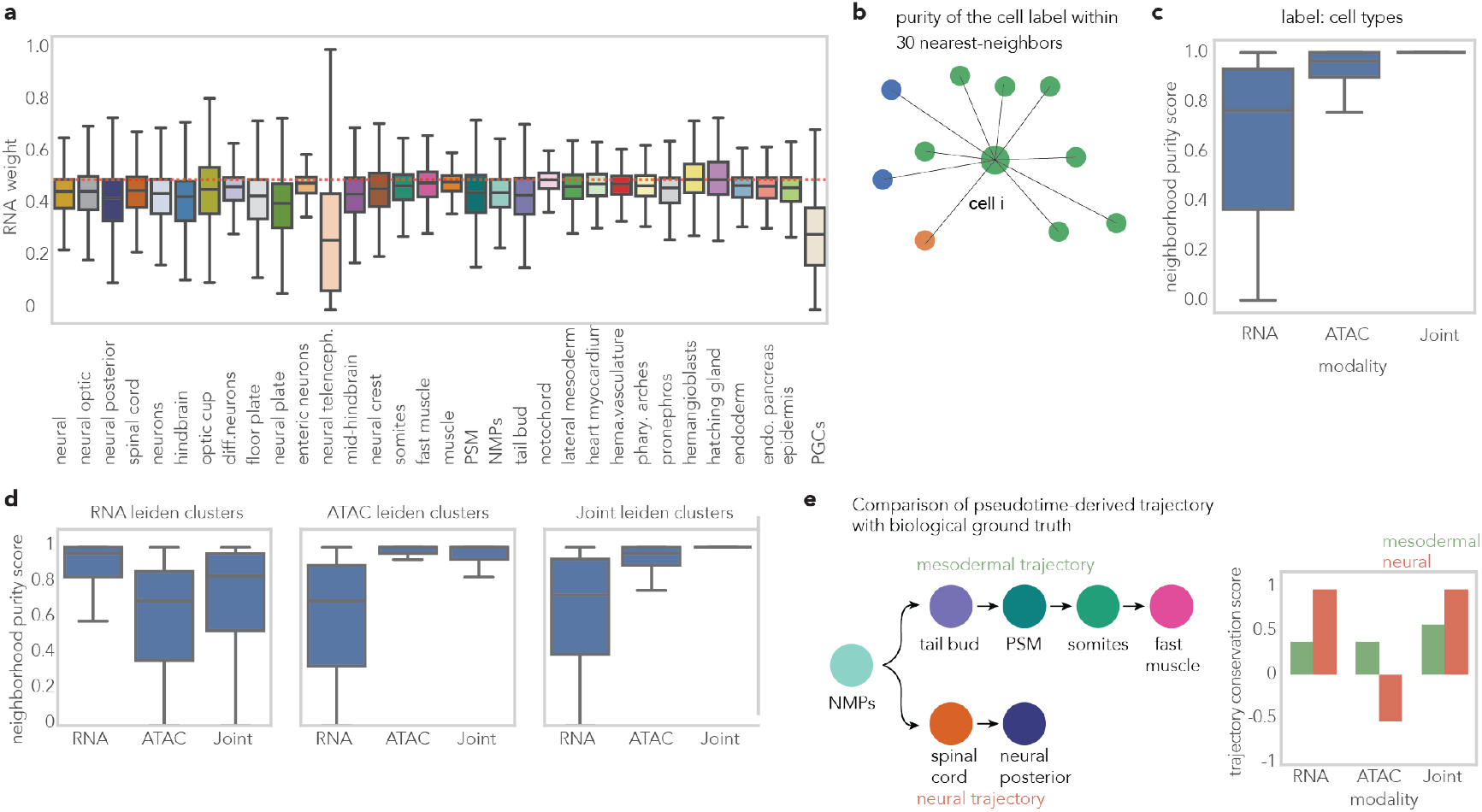
Comparison of neighborhood graphs from RNA, ATAC, and Joint modalities. **a**, Distribution of RNA weights in joint embedding across cell types, showing systematic ATAC dominance (mean RNA weight = 0.444) with 68.9% of cells prioritizing chromatin accessibility over gene expression. **b**, Schematic of neighborhood purity analysis, measuring the proportion of k-nearest neighbors sharing the same metadata label for each cell. **c**, Neighborhood purity for cell type label for each modality. **d**, Neighborhood purity scores for different clustering approaches. Each subplot shows how well RNA, ATAC, and joint neighborhood graphs preserve clusters derived from the corresponding modality, demonstrating that joint neighborhoods achieve high purity across all clustering types. **e**, Trajectory conservation analysis for developmental lineages. Trajectories inferred from diffusion pseudotime are compared with biological ground truth: mesodermal and neural trajectories. The trajectory conservation scales from 0 (no conservation) to 1 (perfect conservation), showing the trajectory conservation for RNA, ATAC, and Joint neighborhood graphs for either mesodermal (green) and neural (red) trajectories.

**Figure S4:**
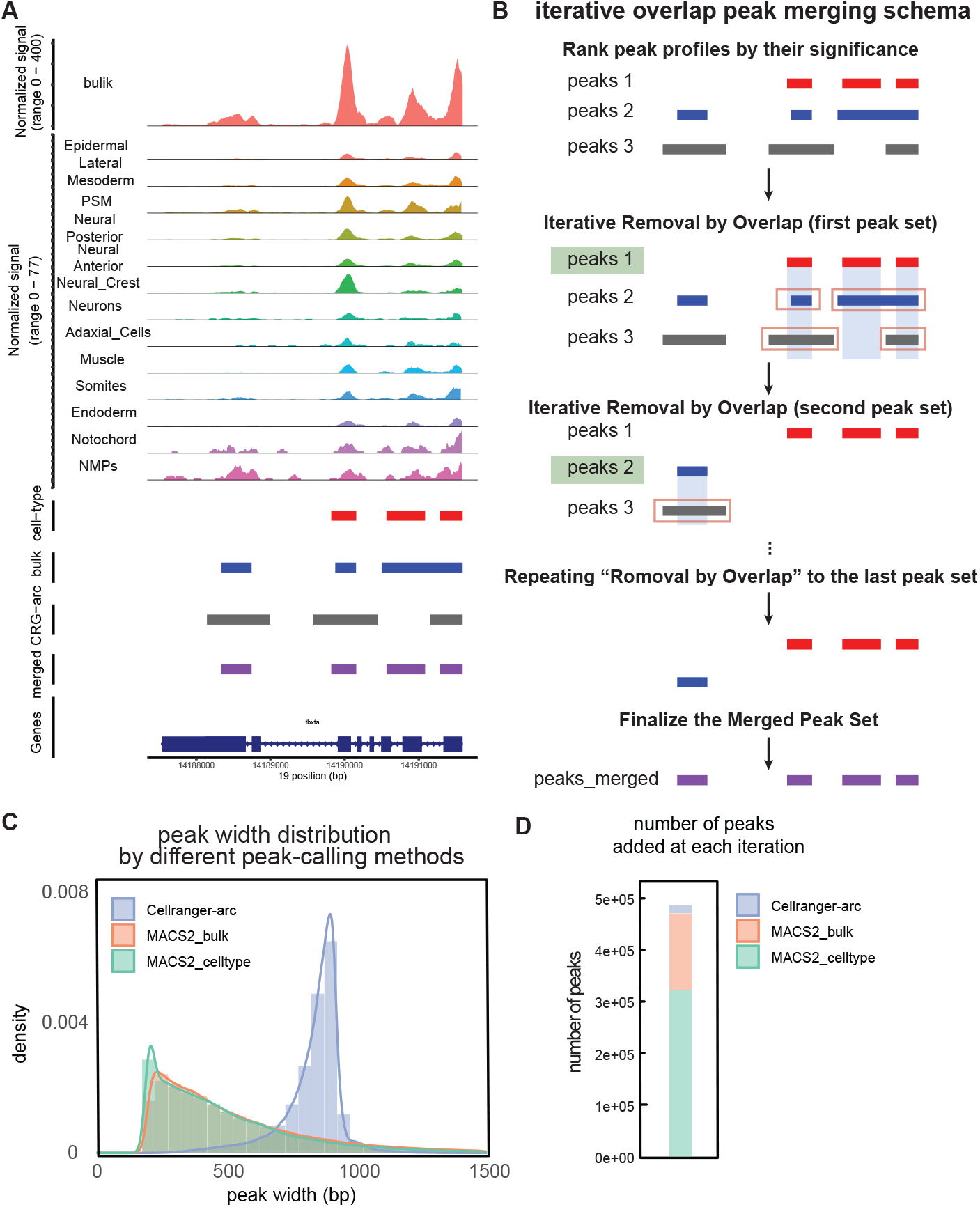
Peak calling strategy for scATAC-seq datasets. **a**, An example genomic coverage plot for *tbxta* gene. The normalized signal on the y-axis represents the pseudo-bulk counts for *tbxta* gene, normalized by the total counts from the specific group of cells. Here, the pseudobulk groups are shown either as bulk (comprising all cells) or as specific cell types, respectively. Four peak profiles are shown below the coverage plot - (1) by MACS2 for each cell type (red), (2) by MACS2 for bulk (blue), (3) by Cellranger-arc for bulk (grey), and (4) by iteratively merging the overlaps for three aforementioned peak profiles. **b**, Iterative overlap peak merging schematics. Peak profiles from different peak-calling metrics were first ranked by their significance. Then, less-significant peaks that overlap with more significant peaks are removed. Lastly, all the remaining peaks are finalized as the unique peak set. **c**, Histogram for the peak width from different peak-calling methods. **d**, Number of peaks added at each iteration in **b**.

**Figure S5:**
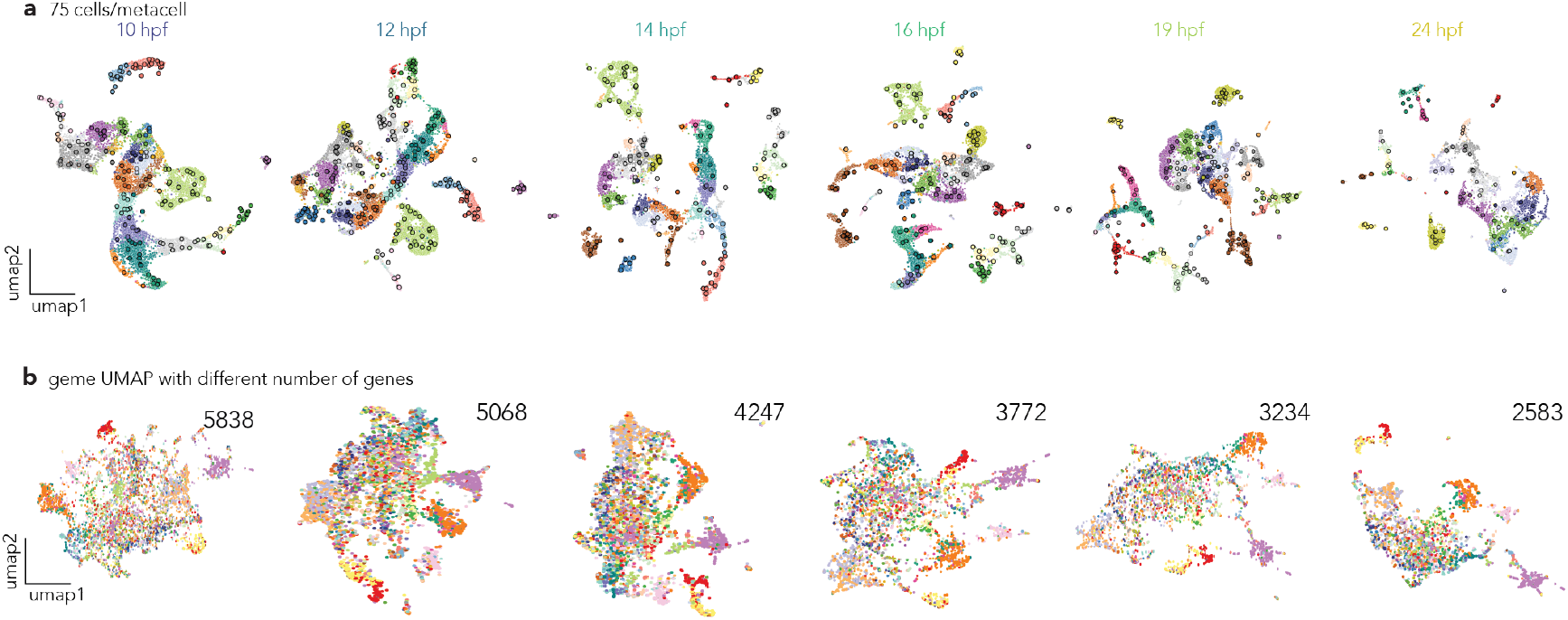
Gene UMAP computed from the metacell aggregation strategy. **a**, A series of cell UMAPs overlaid with metacells for each time point over six developmental stages, metacells computed with 75 cells/metacell as the default parameters. **b**, Gene UMAP computed across varying numbers of highly variable genes (5838 to 2583 genes), where each point represents a gene positioned according to its regulatory dynamics similarity across cell types and developmental stages.

**Figure S6:**
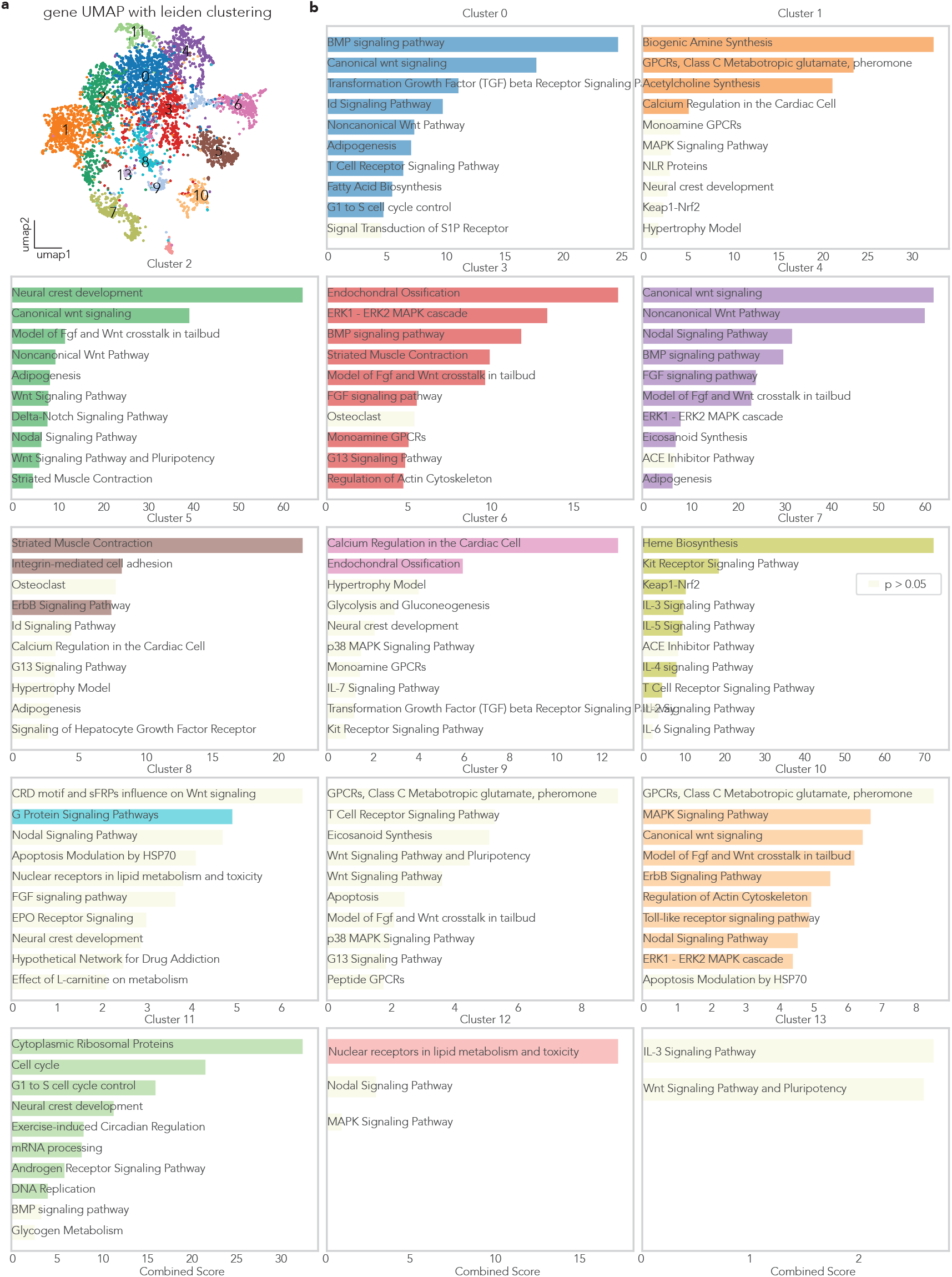
Gene Set Enrichment Analysis for Gene UMAP Leiden clusters. **a**, Gene UMAP colored by 15 distinct leiden clusters. **b**, Gene set enrichment analysis using FishEnrichr for each Leiden cluster, displaying the top 10 significantly enriched biological pathways and processes, with insignificant enrichment is colored beige. Each cluster captures functionally related genes with coordinated expression and accessibility patterns, including early developmental signaling (BMP, Wnt, Nodal signaling in clusters 0, 2, 4, 7, 12), cell cycle regulation (cluster 10), neural development (clusters 2, 5, 6), metabolic processes (clusters 1, 9, 13), and tissue-specific differentiation programs (clusters 3, 6, 8, 11).

**Figure S7:**
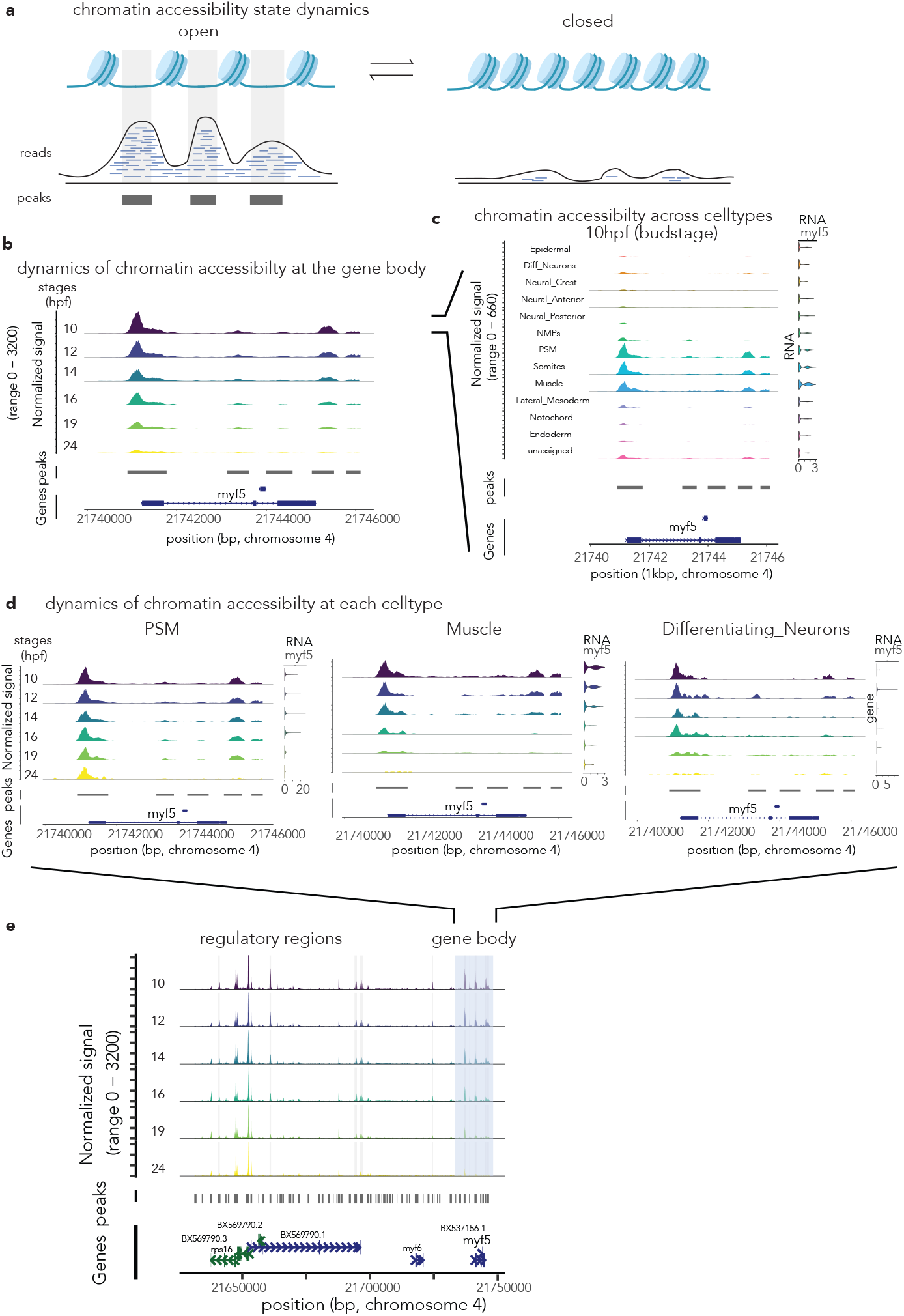
Dynamics of chromatin accessibility during zebrafish development. **a** Schematic representation of chromatin accessibility states and their corresponding ATAC-seq signal profiles. **b** Chromatin accessibility dynamics at the *myf5* gene locus across developmental stages (10 to 24 hpf). **c** Cell-type-specific chromatin accessibility patterns at the *myf5* locus at 10 hpf (budstage). **d** Temporal dynamics of chromatin accessibility at the *myf5* locus in specific cell types: presomitic mesoderm (PSM), muscle, and differentiating neurons. **e** Detailed view of chromatin accessibility changes at the *myf5* locus and surrounding regulatory regions across developmental stages (10 to 24 hpf).

**Figure S8:**
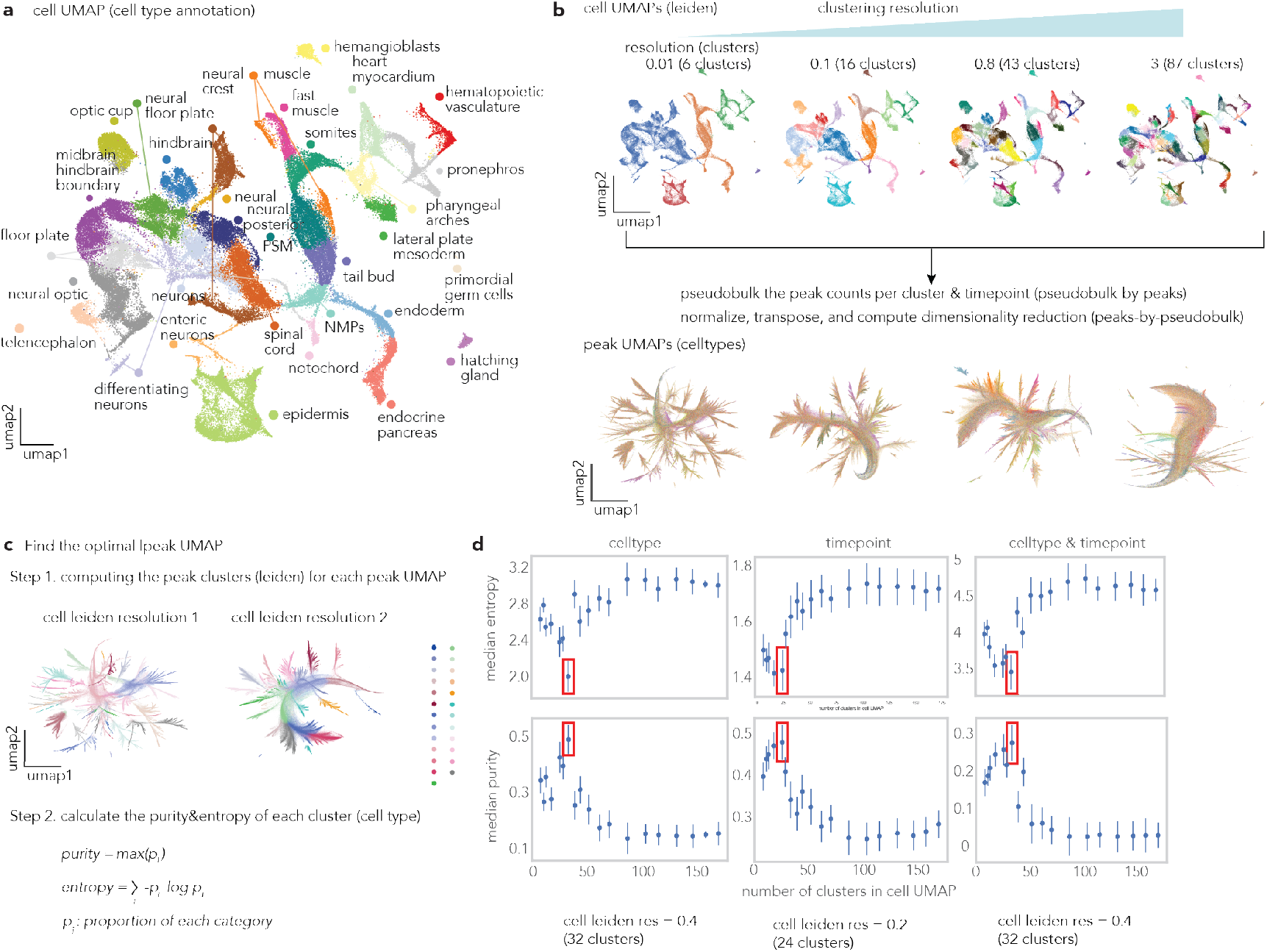
Parameter optimization for pseudobulk strategy through cell clustering resolution assessment. **a**, Cell UMAP colored by cell type annotation in **??** used as input for pseudobulk strategy. **b**, Effect of cell leiden clustering resolution on peak UMAP structure. Different cell clustering resolutions (0.01 with 6 clusters, 0.1 with 16 clusters, 0.8 with 43 clusters, and 3.0 with 87 clusters) were tested to determine the optimal value for pseudobulk resolution. Each cell clustering resolution yields corresponding peak UMAPs (bottom row), where each point represents a chromatin accessibility peak positioned according to its cell type and temporal accessibility patterns. **c**, Optimization workflow schematic showing the two-step process: Step 1 involves computing Leiden clusters for each cell clustering resolution, and Step 2 calculates purity and entropy metrics for each peak cluster to assess cell-type and temporal specificity. **d**, Optimization curves showing median entropy (top row) and median purity (bottom row) across different numbers of cell clusters for three evaluation criteria: cell type specificity, temporal specificity, and combined cell type & temporal specificity. Red boxes indicate optimal cell clustering resolutions (Leiden resolution = 0.4, corresponding to 32 clusters) that maximize peak cluster purity while maintaining sufficient granularity for cell-type and temporal resolution. This systematic optimization ensures that downstream peak clustering analysis captures biologically meaningful chromatin accessibility patterns with optimal cell-type and temporal resolution. **e**, A peak UMAP colored by the most accessible cell type. **f**, A peak UMAP colored by the most accessible time point.

**Figure S9:**
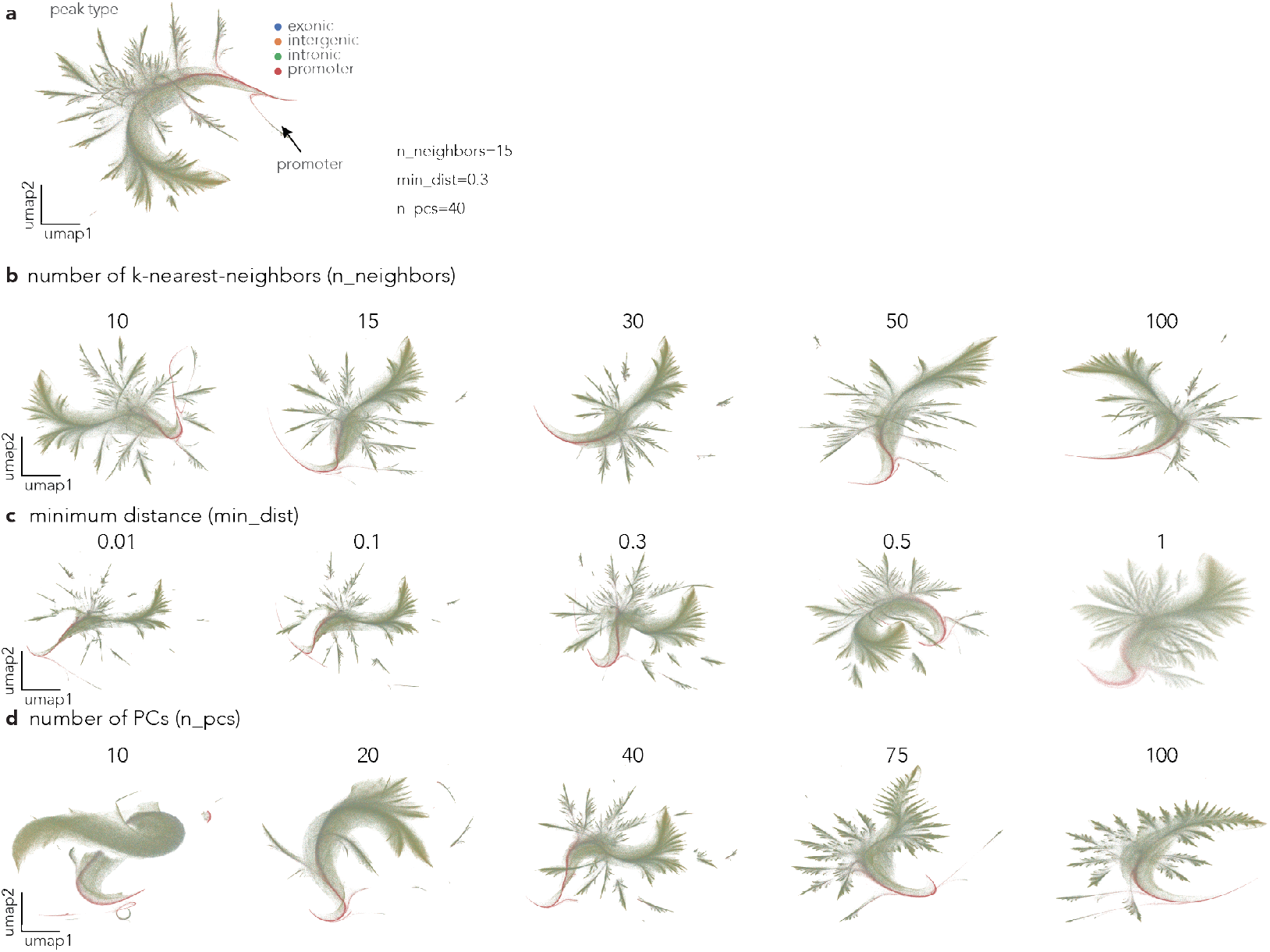
Hyper-parameter exploration for peak UMAP visualization. **a**, Reference peak UMAP computed with default parameters (n neighbors=15, min dist=0.3, n pcs=40) showing the distribution of chromatin accessibility peaks colored by peak type based on genomic annotation. **b**, UMAP visualization with differnt number of k-nearest neighbors (n neighbors) from 10 to 100. **c**, UMAP visualization with different minimum distance parameter (min dist) ranging from 0.01 to 1.0. **d**, UMAP visualization with different number of principal components (n pcs) from 10 to 100

**Figure S10:**
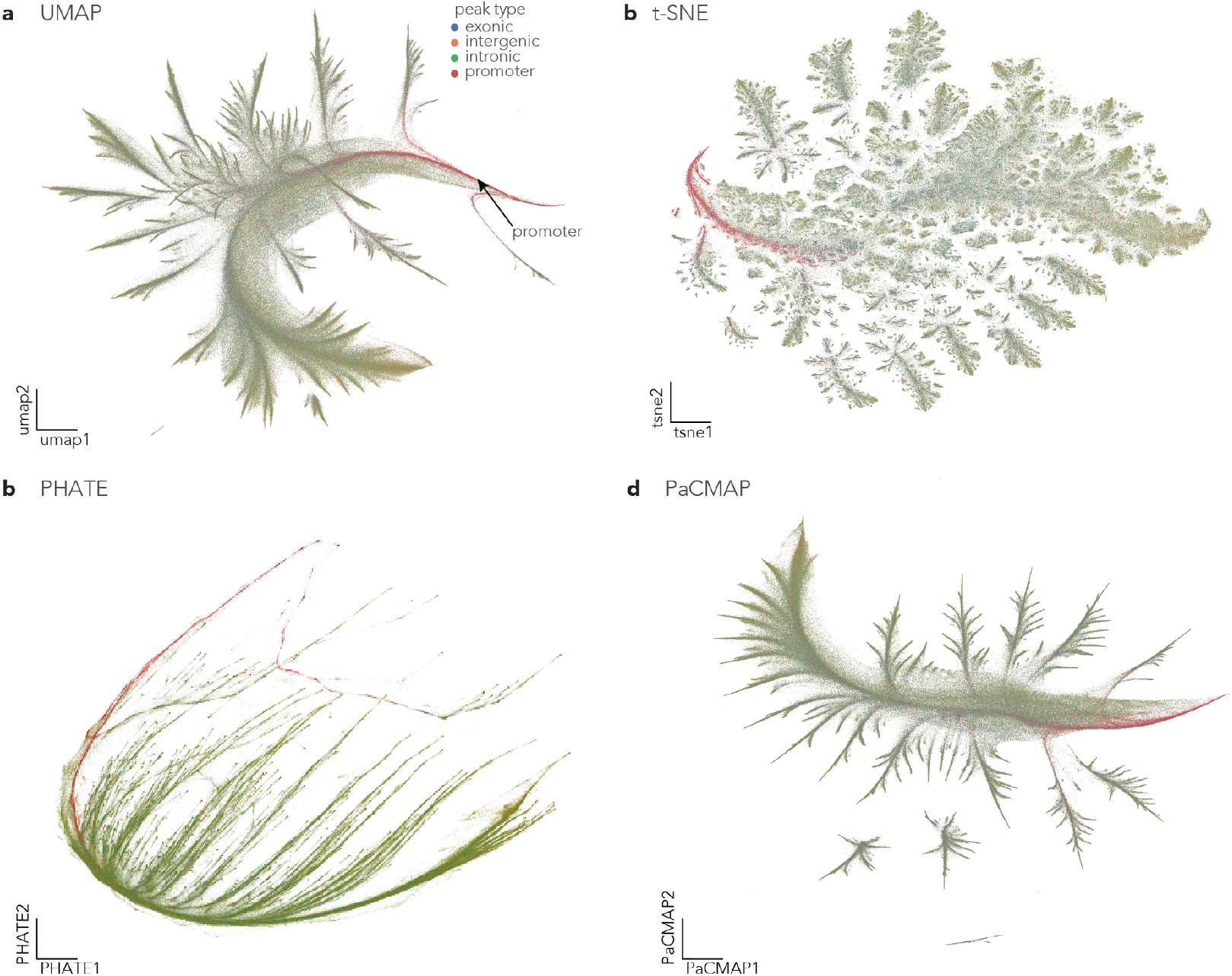
Different dimensionality reduction methods for 2D projection of peaks-by-pseudobulk object. **a**, Reference peak UMAP computed with default parameters (n neighbors=15, min dist=0.3, n pcs=40) showing the distribution of chromatin accessibility peaks colored by peak type based on genomic annotation. **b**, t-SNE^94^ visualization with default parameters. **c**, PHATE^95^ visualization with default parameters. **d**, PaCMAP^96^ visualization with default parameters.

**Figure S11:**
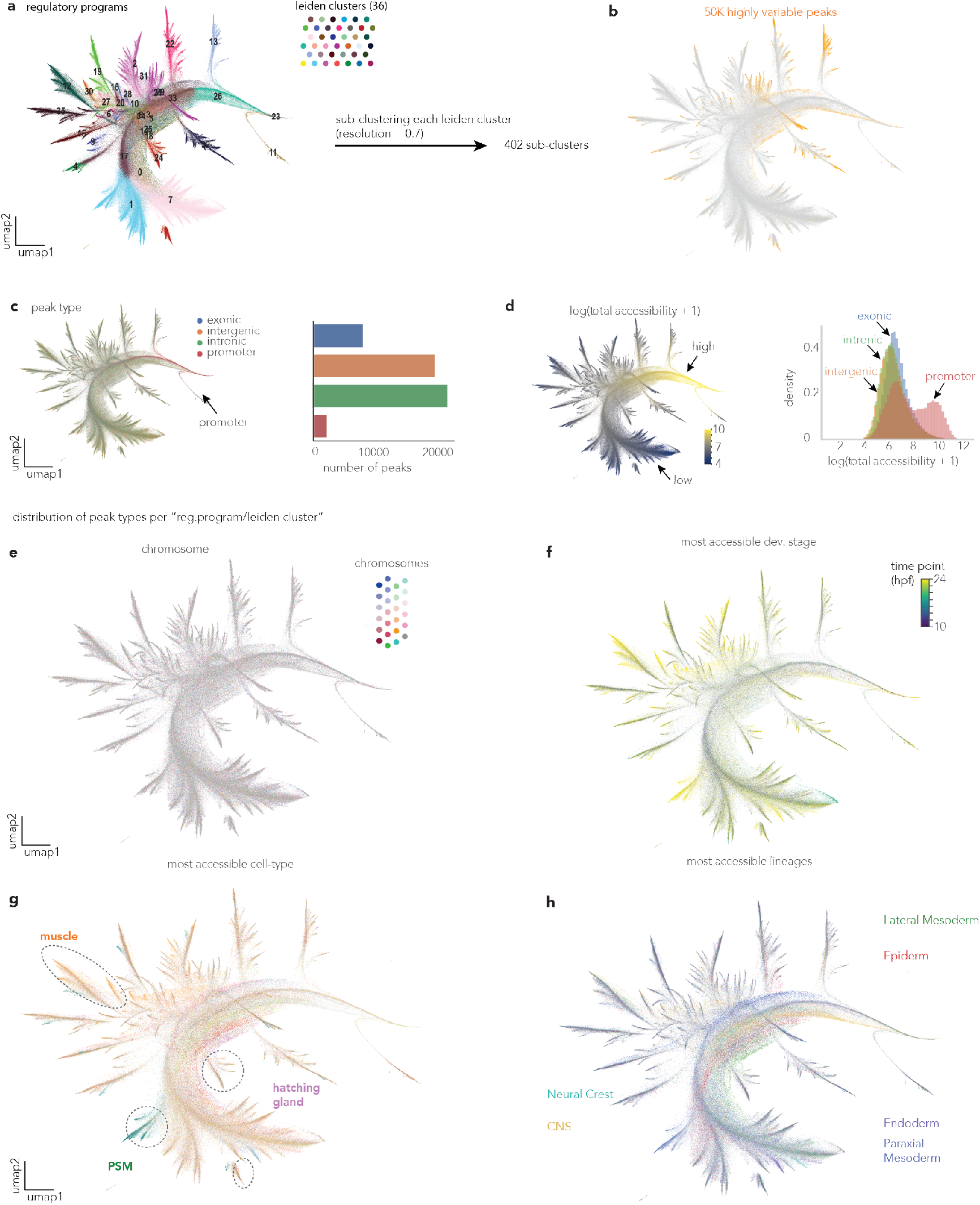
Comprehensive annotation and characterization of genomic regions(peaks) reveals functional organization principles. **a**, Peak UMAP colored by 36 major Leiden clusters (left) with hierarchical sub-clustering generating 402 fine-grained clusters (right). Each point represents a single regulatory element (genomic loci), positioned according to its accessibility dynamics across different cell types and developmental stages. **b**, Identification of 50,000 highly variable peaks (highlighted in orange) from the complete dataset of 640,830 regulatory elements (grey) for downstream analysis. **c**, Peak type annotation showing genomic distribution across exonic, intergenic, intronic, and promoter regions. A bar plot quantifies the number of peaks per category. Promoter peaks (red) cluster distinctly at the root of the UMAP (indicated by the arrow). **d**, Peak accessibility levels visualized as total accessibility scores summed over all pseudobulk groups, with histogram showing the distribution. **e**, Peak UMAP colored by chromosomal location, demonstrating lack of chromosome-specific clustering patterns in the accessibility landscape. **f**, Temporal dynamics of peak accessibility colored by the developmental stage showing maximum activity (10-24 hpf), revealing stage-specific regulatory programs. **g**, Cell-type specificity analysis showing peaks colored by their most accessible cell type, with specific examples highlighted, including muscle, hatching gland, and PSM (presomitic mesoderm) regulatory programs. **h**, Lineage-specific accessibility patterns colored by major developmental lineages (Lateral Mesoderm, Epiderm, Neural Crest, CNS, Endoderm, Paraxial Mesoderm), demonstrating coordinated regulatory programs within related cell types.

**Figure S12:**
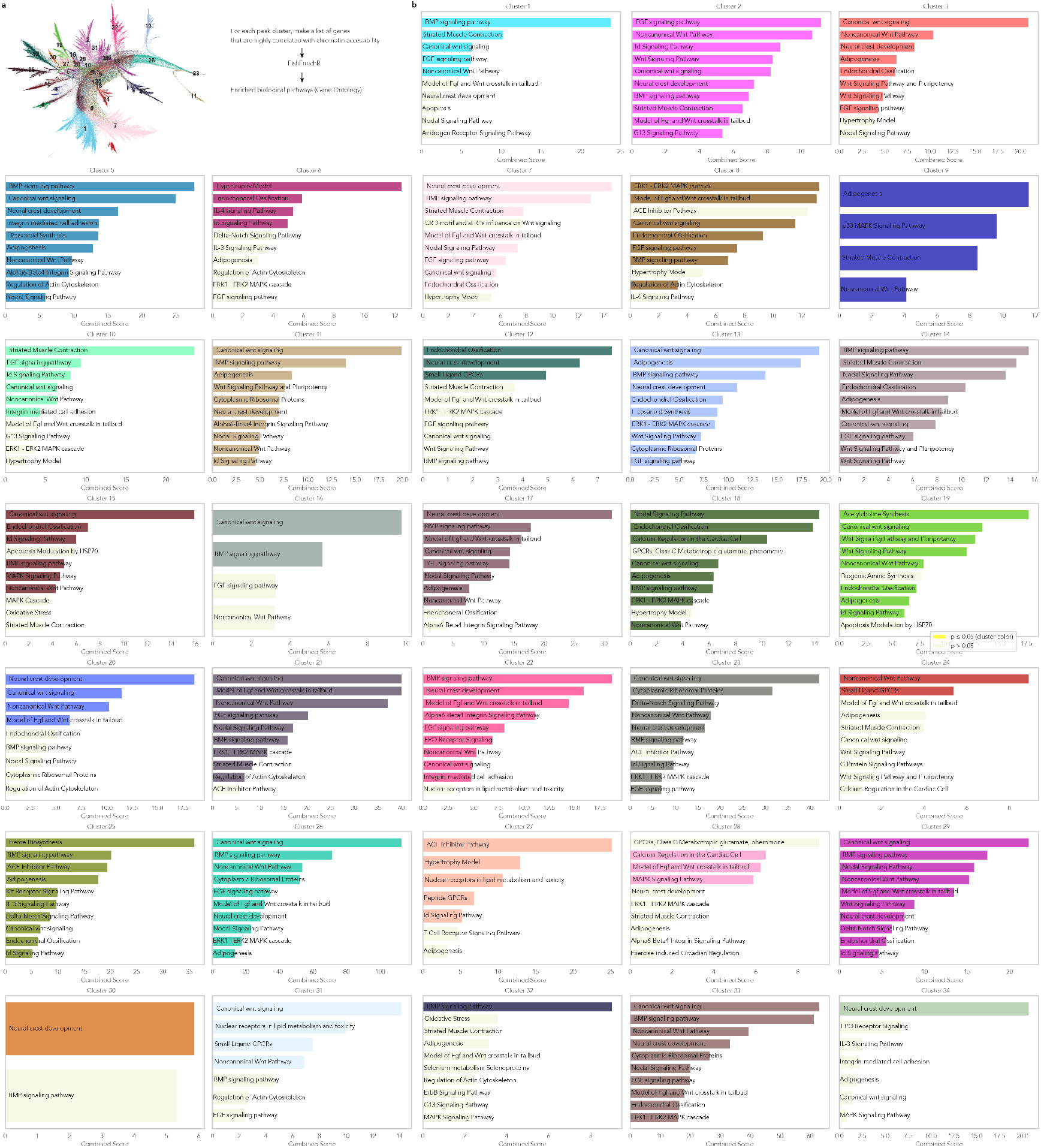
Functional annotation of peak cluster-associated genes reveals cell-type and stage-specific regulatory programs. **a**, Peak UMAP with Leiden clustering and workflow for gene set enrichment analysis using FishEnrichR. For each peak cluster, a list of genes highly correlated with peak accessibility patterns was identified and subjected to functional enrichment analysis using the zebrafish-specific FishEnrichR database. **b**, Gene set enrichment analysis results using FishEnrichR for each Leiden peak cluster, displaying the top significantly enriched biological pathways and processes using the genes associated with the peaks from each cluster. Peak clusters capture functionally coherent regulatory programs, including early developmental signaling (BMP, Wnt, Nodal signaling pathways in multiple clusters), neural development (neural crest development, canonical Wnt signaling), muscle differentiation (striated muscle contraction, ACE inhibitor pathway), metabolic processes (heme biosynthesis, lipid metabolism), and tissue-specific differentiation programs across the 36 identified peak clusters.

**Figure S13:**
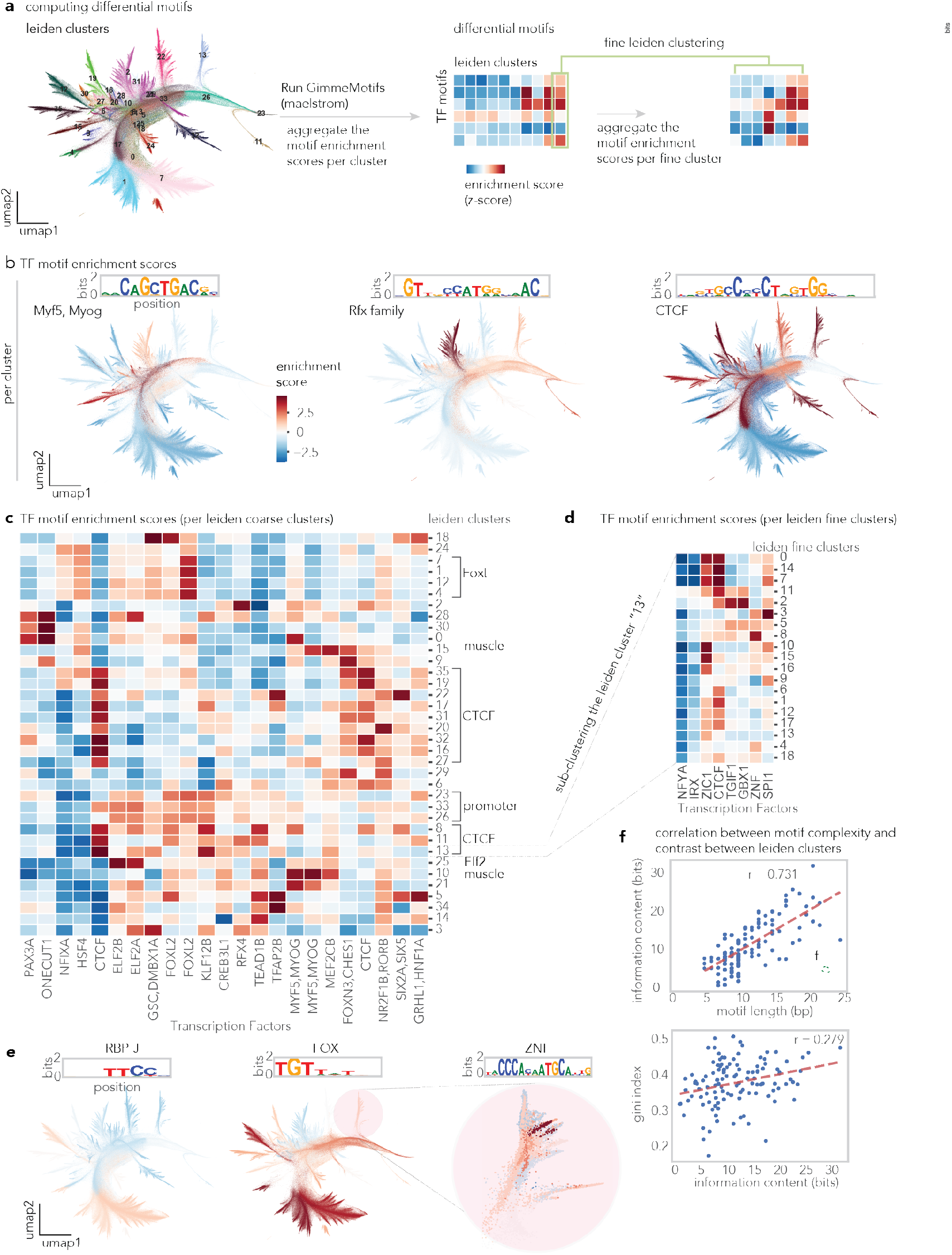
Transcription factor motif enrichment analysis for the peak UMAP. **a**, Peak UMAP with Leiden clustering and workflow for differential motif enrichment analysis using GimmeMotifs Maelstrom to identify transcription factor binding motifs enriched within specific peak clusters. **b**, Representative transcription factor motif families showing sequence logos and corresponding peak UMAP colored by motif enrichment scores (z-scores). Examples include Myf5/Myog (muscle-specific), the Rfx family (involved in cilia and neural tube development), and CTCF (involved in chromatin organization). **c**, Comprehensive heatmap of transcription factor motif enrichment scores across Leiden coarse clusters (rows) and transcription factors (columns), revealing cluster-specific regulatory signatures. Notable enrichments include FoxL and muscle-related factors in specific clusters, promoter-associated factors, and CTCF binding patterns. **d**, Fine-resolution transcription factor motif enrichment analysis using Leiden fine clusters, showing more granular regulatory program identification with enhanced resolution of cell-type and temporal specificity. **e**, Individual motif examples (RBP-J, FOX, ZNF families) displaying sequence logos and peak UMAP visualization colored by motif enrichment, demonstrating cluster-specific transcription factor binding patterns. **f**, Correlation between motif complexity measures: motif length correlation with information content (r = 0.731) and information content correlation with Gini index (r = 0.279).

**Figure S14:**
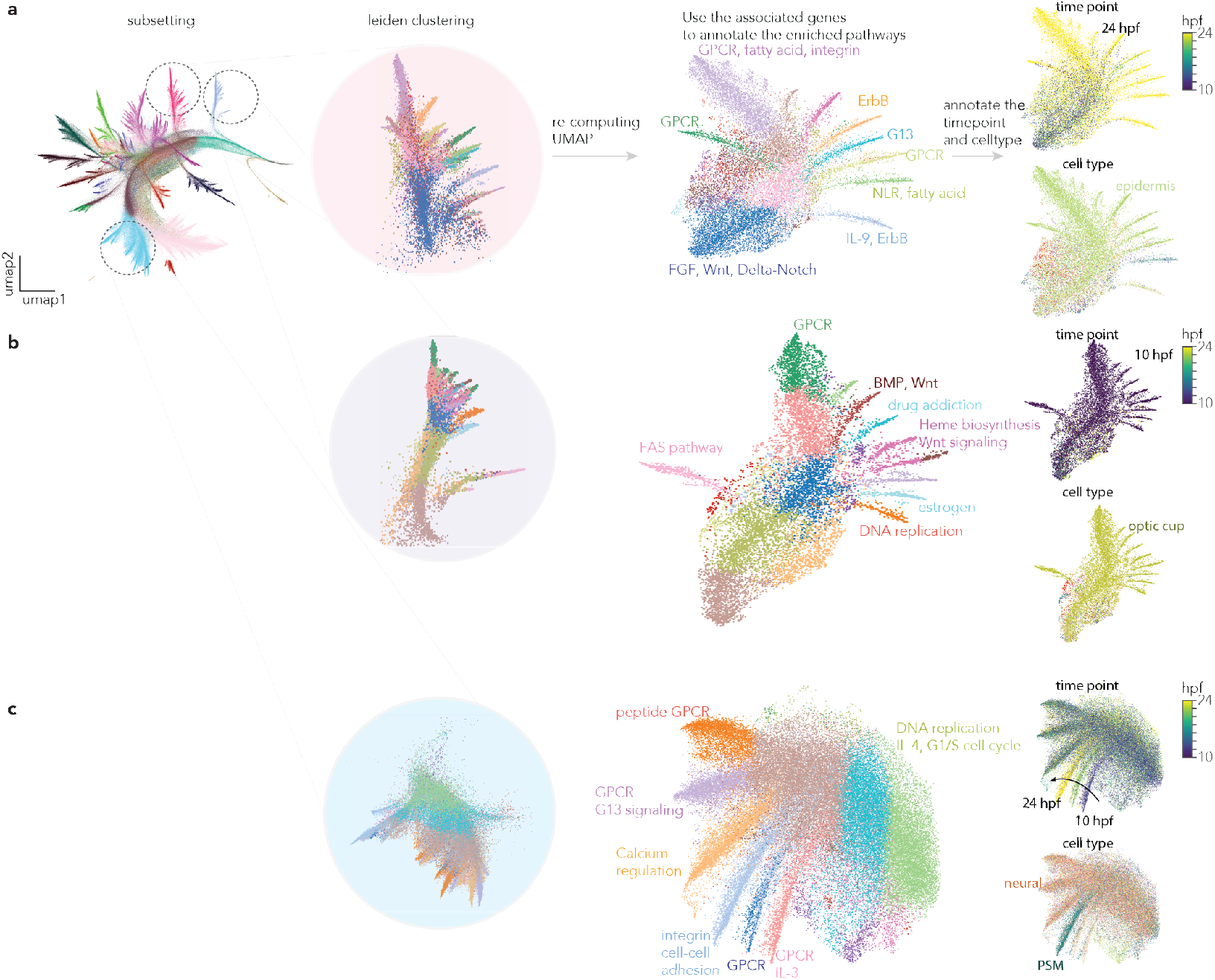
Peak UMAP sub-clustering and annotation of enriched biological pathways. **a**, Peak UMAP sub-clustering scheme (left). Coarse peak cluster 22 (left) is subsetted and Leiden clustering is performed, and visualized with re-embedding. Recomputed UMAP for each coarse cluster, colored by sub-clusters, is highlighted with enriched biological pathways ^43^(center), and the most enriched time point and cell type (right). **b, c** Re-computed UMAPs colored by sub-clusters and enriched biological pathways (center), enriched cell type and time point (right) for coarse cluster **b**, 13, and **c**, 1.

**Figure S15:**
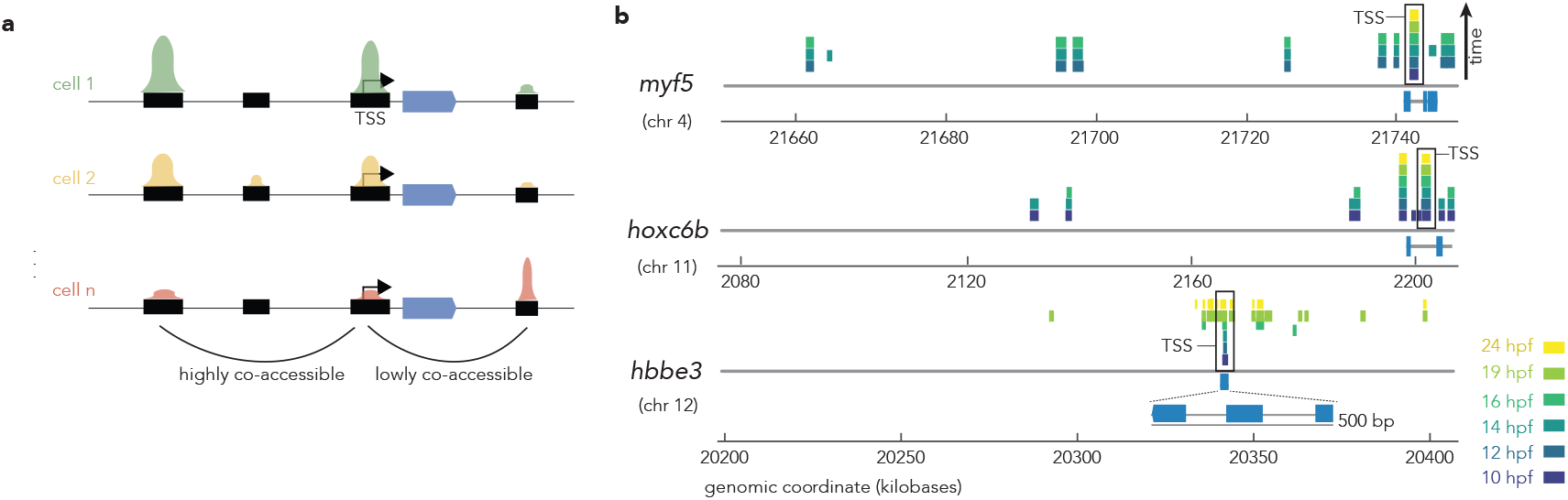
Temporal dynamics of chromatin co-accessibility. **a**, A schematic illustrating chromatin co-accessibility. **b**, Highly co-accessible peaks with transcription start site for *myf5, hoxc6b*, and *hbbe3* for each developmental stage along the genomic tracks.

**Figure S16:**
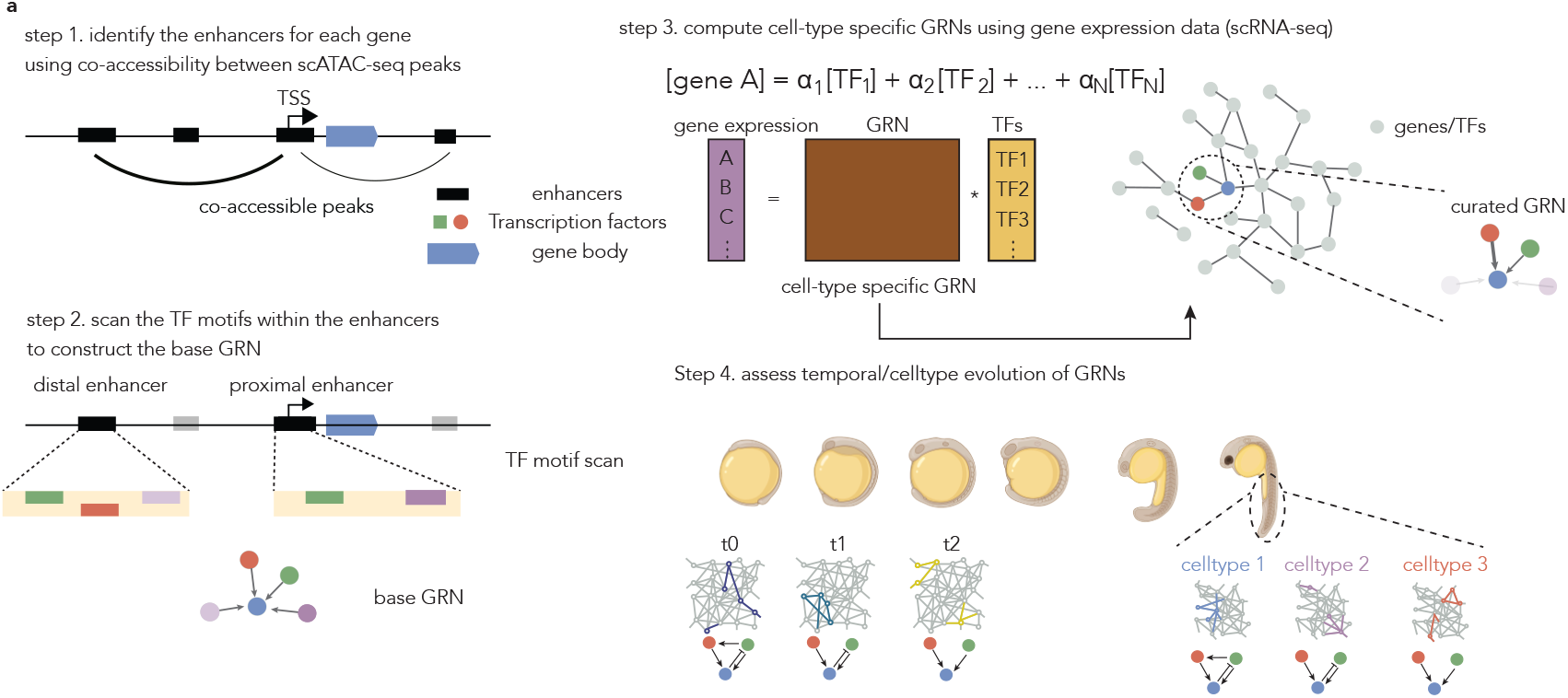
CellOracle workflow schematics. a, A schematic of CellOracle workflow to compute the gene regulatory network.

**Figure S17:**
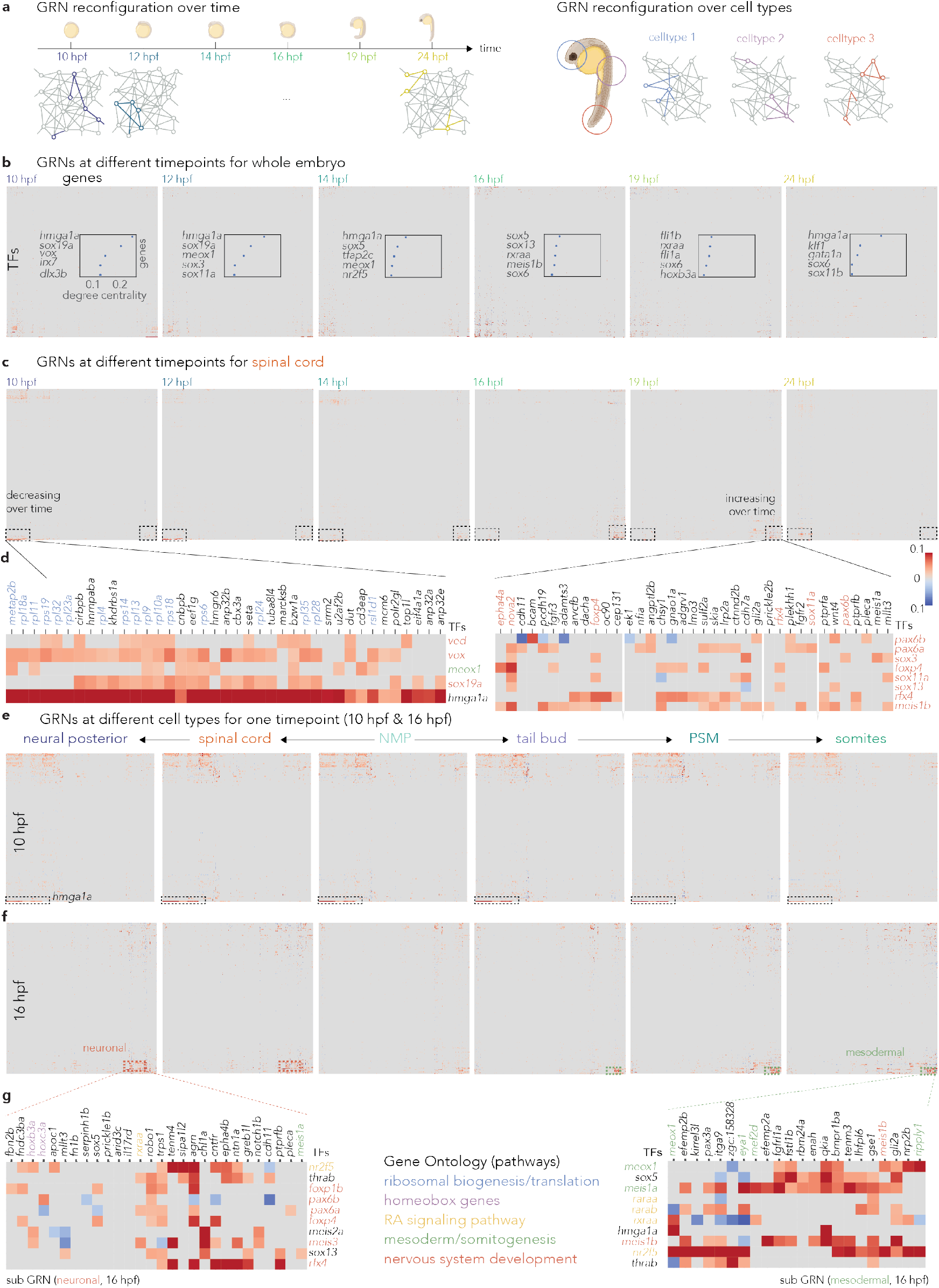
Dynamic Gene Regulatory Networks (GRNs) across developmental stages and cell types in zebrafish embryogenesis. **a**, A schematic representation of GRN reconfiguration (1) over developmental time and (2) across different cell types. **b**, GRNs for whole embryo across developmental stages (10 to 24 hpf). Heatmaps display the edge strengths between transcription factors (TFs) and their target genes. The top five transcription factors with the highest degree centrality at each time point are shown with their degree centrality score. **c**, GRNs for spinal cord development across different developmental stages (10 to 24 hpf). Heatmaps display the edge strengths between transcription factors (TFs) and target genes, with both decreasing and increasing patterns highlighted. **d**, Detailed view of the GRN heatmaps for spinal cord, showing specific TF-gene interactions. Edge strength is represented by color intensity, with red indicating positive and blue indicating negative regulation. **e, f**, GRNs for different cell types at the (e) 10 hpf, and (f) 24 hpf stages, including Neural Posterior, Spinal Cord, Neuromesodermal Progenitors (NMP), Tail Bud, Presomitic Mesoderm (PSM), and Somites. **g**, Sub-GRNs of the 15-somite stage GRN for neuronal (left) and mesodermal (right) lineages. Gene Ontology pathways associated with specific modules are annotated. Notable TFs and their target genes are highlighted, demonstrating lineage-specific regulatory patterns.

**Figure S18:**
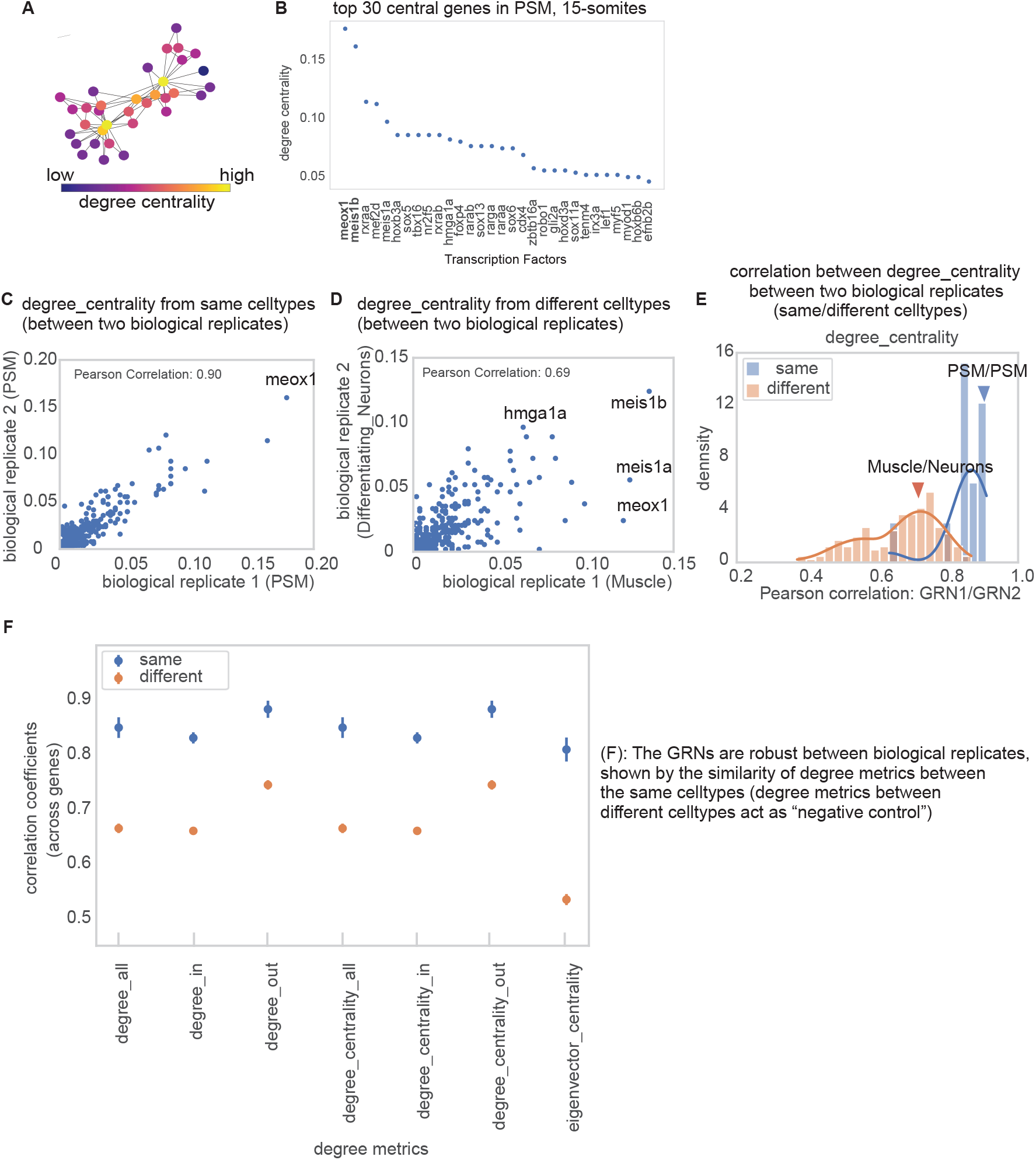
Comparison of GRNs between biological replicates. **a**, An example network diagram with nodes colored by the degree centrality. **b**, Top 30-degree central genes from an example group-PSM (pre-somitic mesoderm) at 16 hpf. **c**, A scatter plot showing the degree centrality of each gene from two GRNs from biological replicates. For example, GRNs from PSM are selected. **d**, A scatter plot showing the degree centrality of each gene from two GRNs from biological replicates - as a negative control, the GRNs from two different cell types were chosen. **e**, A distribution of correlation coefficients of degree centrality from two GRNs between the same cell type (blue), and different cell types (orange). Some example cell type pairs are highlighted. **f**, Correlation coefficients of degree metrics from two GRNs between same (blue) and different (orange) cell types, computed for different degree metrics - degree all, degree in, degree out, degree centrality in, degree centrality out, eigenvector centrality.

**Figure S19:**
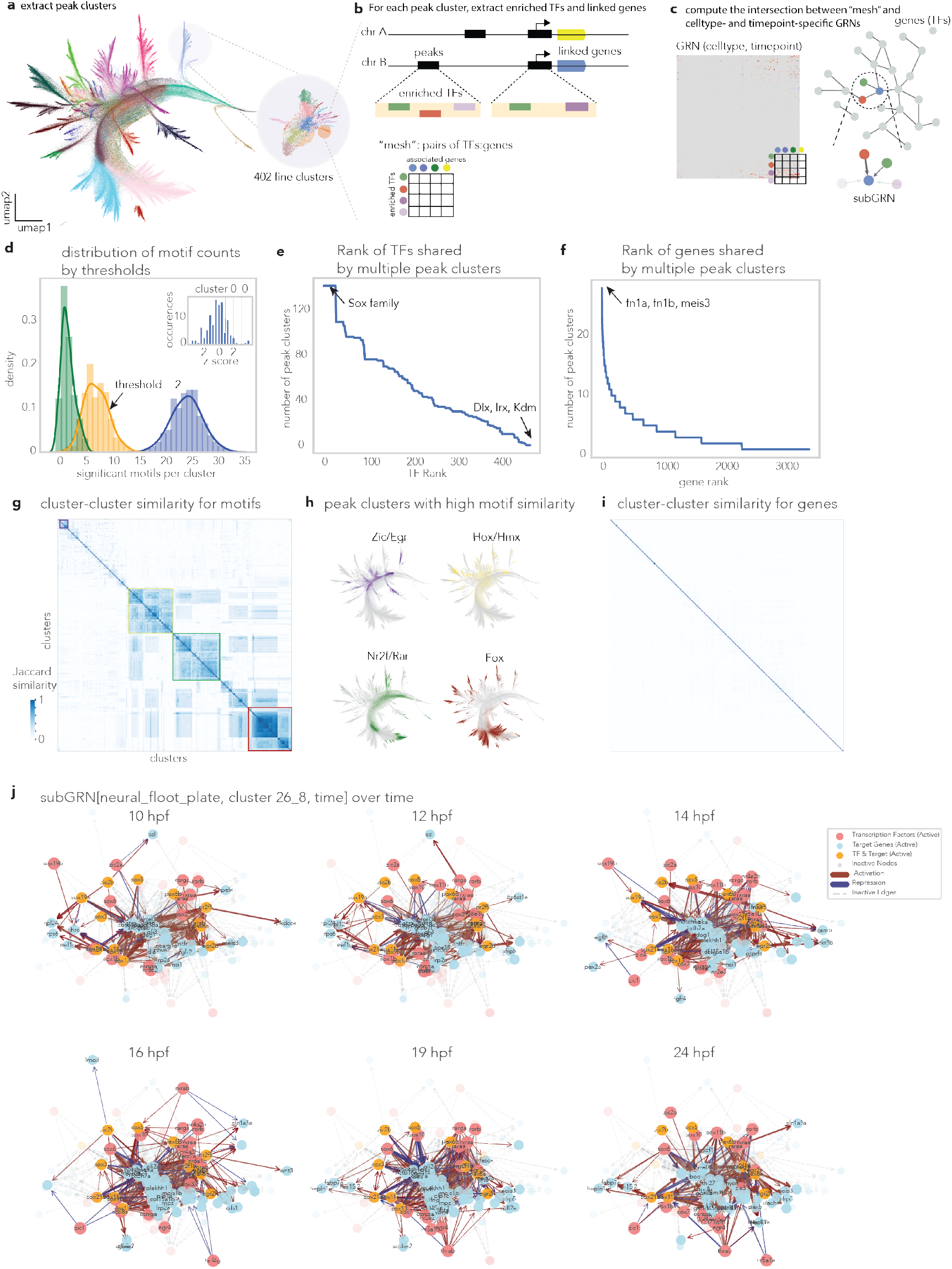
Extraction of subGRNs from celltype- and timepoint-specific GRNs from CellOracle using peak-cluster-specific regulatory programs. **a**, Peak UMAP showing extraction of 402 fine-grained clusters for systematic regulatory program characterization. **b**, Schematic workflow for creating regulatory program “meshes.” For each peak cluster, enriched transcription factors (identified through motif analysis) and linked genes (determined by correlation-based analysis) are identified to generate binary TF-by-gene matrices. **c**, Integration approach combining regulatory program meshes with cell-type and time-point-specific GRNs from CellOracle to extract biologically relevant subGRNs. **d**, Distribution of significant motifs per cluster across different threshold values, with threshold = 2 (arrow) selected to balance sensitivity and specificity. **e**, Ranked distribution showing transcription factors shared across multiple peak clusters, with Sox family members and Dlx/Irx/Kdm factors highlighted as broadly active regulators. **f**, Ranked distribution of genes associated with multiple peak clusters, with Fn1a/Fn1b/Meis3 representing highly connected regulatory targets. **g**, Cluster-cluster similarity matrix for transcription factor motifs, revealing distinct regulatory signatures across peak clusters. **h**, Examples of peak clusters with high motif similarity, showing specific transcription factor families (Zic/Egr, Hox/Hmx, Nr2f/Rar, Fox) that define related regulatory programs. **i**, Cluster-cluster similarity matrix for associated genes, demonstrating functional coherence within regulatory modules. **j**, Temporal dynamics of an example subGRN extracted from neural floor plate cells using regulatory program cluster 26-8. Network topology illustrates transcription factors (red), target genes (blue), and dual-function nodes (yellow) across various developmental time points (from 10 hpf to 24 hpf), with active edges in color and inactive edges in gray, highlighting the regulatory circuit rewiring that occurs during development.

